# Plant-derived benzoxazinoids act as antibiotics and shape bacterial communities

**DOI:** 10.1101/2021.01.12.425818

**Authors:** Niklas Schandry, Katharina Jandrasits, Ruben Garrido-Oter, Claude Becker

## Abstract

Plants synthesize and release specialized metabolites into their environment that can serve as chemical cues for other organisms. Metabolites that are released from the roots are important factors in determining which microorganisms will colonize the root and become part of the plant rhizosphere. Root exudates can be converted by soil microorganisms, which can result in the formation of toxic compounds. How individual members of the plant rhizosphere respond to individual compounds and how the differential response of individual microorganisms contributes to the response of a microbial community remains an open question. Here, we investigated the impact of derivatives of benzoxazinoids, a class of plant root exudates released by important crops such as wheat and maize, on a collection of 180 root-associated bacteria. Phenoxazine, derived in soil from benzoxazinoids, inhibited the growth of root-associated bacteria *in vitro* in an isolate-specific manner, with sensitive and resistant isolates present in most of the studied clades. Using synthetic communities, we show that community stability is a consequence of the resilience of its individual members, with communities assembled from tolerant isolates being overall more tolerant to benzoxazinoids. However, the performance of an isolate in a community context was not correlated with its individual performance but appeared to be shaped by interactions between isolates. These interactions were independent of the overall community composition and were strain-specific, with interactions between different representatives of the same bacterial genera accounting for differential community composition.

## Introduction

In the rhizosphere, the space directly surrounding the roots, plants recruit and maintain a diverse community of microorganisms that differs from the surrounding bulk soil (Bulgarelli et al., 2013, 2012; Fitzpatrick et al., 2020; Stringlis et al., 2018). These rhizospheric microbiota, in particular bacteria and fungi, contribute to plant nutrition, growth, and defense (Fitzpatrick et al., 2020). Recent studies have shown that some plant specialized metabolites contribute to plant species-specific community profiles in the rhizosphere. For example, salicylic acid, camalexin, coumarins, and glucosinolates play an important role in shaping the *Arabidopsis thaliana* rhizosphere (Harbort et al., 2020; Jacoby et al., 2020; Koprivova et al., 2019; Lebeis et al., 2015; Stringlis et al., 2018; Voges et al., 2019).

Members of the Poaceae family, including wheat and maize, produce a particular class of specialized metabolites, the benzoxazinoids (BX) (Frey et al., 1997; Handrick et al., 2016). BX exert a major influence on root-associated fungal and bacterial microbiota. For example, BX exudation by maize plants into the surrounding soil conditions the microbiome, which in turn results in soil-memory effects that affect plants of the following sowing (Hu et al., 2018). Other studies showed that BX exudation specifically recruits beneficial pseudomonads to the rhizosphere, whereas *Comamonadaceae* are often depleted by BX exudation (Cadot et al., 2020; Cotton et al., 2019; Kudjordjie et al., 2019; Neal et al., 2012; Schandry and Becker, 2020). Conversely, the presence and composition of the microbiota can alter BX chemistry. Post-release breakdown and metabolism of benzoxazinoid compounds by soil microbiota leads to the sequential formation of 1,4-benzoxazin-3-one (BOA) and 2-amino-3-phenoxazinone (APO), among other compounds, in non-sterile soils (Kumar et al., 1993; Macías et al., 2005, 2004). APO in return exhibits antibiotic activity similar to other phenoxazines and limits plant root growth at concentrations corresponding to those detected in soil (Anzai et al., 1960; Cheng et al., 2016; Imai et al., 1990; Venturelli et al., 2016, 2015).

Given the long-term microbiome-conditioning capacity of BX exudation (Hu et al., 2018), it is tempting to speculate that some of the plant-growth-inhibitory effects of BX compounds could be mediated by selective effects on the plant-associated microbiota (Belz and Hurle, 2005). However, quantitative analyses of the degree to which microbial isolates differ in their response to root exudates are extremely difficult to perform in soil, because (*i*) plant-associated microbiota are highly diverse, and (*ii*) plant exudates are typically complex mixtures, making it difficult to attribute effects to a particular chemical. In contrast, such studies can be carried out *in vitro* using clonal bacterial populations (isolates) and pure compounds at defined concentrations. Collections of culturable bacterial isolates from diverse habitats are an important resource to investigate how diverse microorganisms respond to chemical exposure (e.g., antibiotics, toxins, or pollutants). For example, studies on the effect of therapeutic drugs on an isolate collection derived from the human gut revealed isolate-specific effects of non-antibiotic therapeutics on bacterial growth (Maier et al., 2018).

Such collections can moreover form the basis for rationally designed synthetic microbial communities (SynComs) (Herrera Paredes et al., 2018; Vorholt et al., 2017). SynComs have been proposed and used as reduced complexity systems to study how various microbiomes assemble and develop over time (De Roy et al., 2014; Finkel et al., 2020; Liu et al., 2019; Voges et al., 2019; Vrancken et al., 2019). Bacterial SynComs in particular have been adopted in plant sciences to study various aspects of plant biology, including plant phenotypes under limiting phosphate conditions (Herrera Paredes et al., 2018), assembly of the phylosphere microbiome (Bodenhausen et al., 2014; Carlström et al., 2019), or assembly cues of the root microbiome (Bai et al., 2015; Finkel et al., 2020; Voges et al., 2019). While host-free *in vitro* systems have already been adopted in other systems, e.g. gut microbiota (Li et al., 2019), SynComs assembled from plant-associated bacteria have not yet been studied for their ability to assemble stable communities *in vitro* in the absence of a host, although such a system would significantly reduce the experimental complexity.

Here, we investigated the impact of BX and their derivatives on bacterial members of the plant root and rhizosphere by exposing these isolates to varying concentrations of APO and BOA, two BX detectable in soil surrounding BX-releasing plants. We used the *AtRSphere* collection of culturable *Arabidopsis thaliana-derived* bacterial isolates, spanning four bacterial phyla (Bai et al., 2015); *A. thaliana* is not a BX producer and hence the isolates’ capacity to colonize its roots should be independent of their BX tolerance. In line with this assumption, growth responses, most notably those to APO, of *AtRSphere* isolates varied substantially both within and across genera, ranging from complete insensitivity to strongly impeded growth.

We designed different SynComs to investigate how isolate-level responses to BX influence *in vitro* community assembly and resilience to BX-mediated community perturbations. An isolate’s individual growth behaviour did not correlate with its ability to establish as a community member or with its sensitivity to BX when growing in a community. Moreover, there was no phylogenetic signal for BX tolerance within most bacterial orders, indicating that BX tolerance is strain-rather than genus- or family-specific. Finally, through comparative analyses of isolates and genera shared between differently composed SynComs, we show that BX-mediated restructuring of bacterial communities is attributable to the selective inhibition of some members that relieves antagonistic interactions with other community members.

## Results

### BX derivatives selectively inhibit the growth of root associated bacteria *in vitro*

To gain an overview of how benzoxazinoids and their derivatives affect individual plant-associated bacteria, we screened the *AtRSphere* collection comprising 180 isolates of *A. thaliana* rhizospheric bacteria (Bai et al., 2015) (Figure 1). We monitored the growth of each isolate over 70 h in half-strength TSB medium after supplementation with either BOA (Figure 1A, inner heatmap) or APO (Figure 1A, outer heatmap), two plant-BX-derived compounds commonly found in soil, and under control conditions (equivalent concentration of solvent only). By determining cell density at regular intervals and comparing the area under the curve for the control and BX treatments, we assessed each isolate’s response to different compound concentrations (Figure 1A).

**Figure 1.**
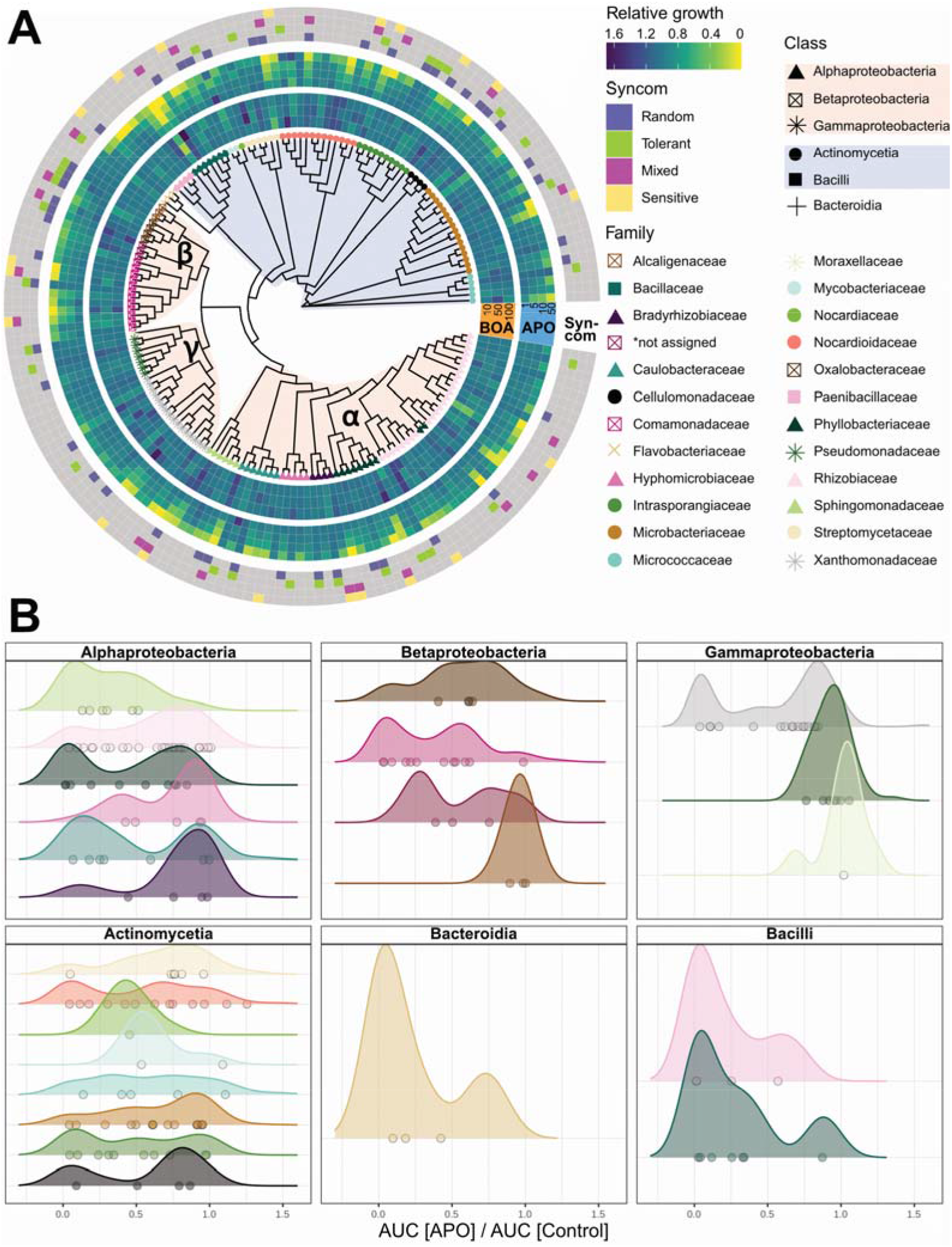
Isolate sensitivity to BX compounds and derivatives. **A)** Heatmap arranged around a circular unrooted phylogenetic tree based on several marker genes. Nodes represent isolates from the collection, node shape indicates taxonomic class, node color indicates taxonomic family (genera Rhizobacter and Methylibium are not assigned to a taxonomic family). The inner heatmap displays average relative growth upon BOA treatment (from inner to outer: 10 μM, 50 μM and 100 μM BOA, rings 1-3). The central heatmap displays relative growth upon APO treatment (inner to outer: 1 μM, 5 μM, 10 μM and 50 μM, rings 4-7). Relative growth was estimated from at least 6 replicate experiments per isolate. The outer heatmap indicates whether or not the isolate was selected as a member of one of the four synthetic communities (see main text for details). **B)** Density plots of relative growth at 50 uM APO. Densities are shown per family (same colors as in A), per class. Dots show the mean relative growth of individual isolates belonging to the respective family.

Because the strains had been isolated from *A. thaliana*, a species that does not release BX, we hypothesized that the collection should cover a broad diversity in terms of BX tolerance. We used 50% relative growth compared to control as the threshold to classify isolates as either sensitive (<50%) or tolerant (>50%).

The growth of most isolates was not inhibited by BOA, even at high concentrations of 100 μM. Some isolates even seemed to benefit (darker color in Figure 1) from the presence of BOA compared with control conditions; this was most consistent for isolates within the *Rhizobium* and *Paenibacillus* genera (Figure 1A). However, such growth promotion was rare overall; this might be attributable to the use of a nutrient-rich medium in this study, which would make BOA superfluous as a potential energy source.

The BX derivative APO was previously shown to have antibiotic effects, so we anticipated a more pronounced negative and dose-dependent effect of APO on isolate growth (Anzai et al., 1960). Indeed, some isolates were insensitive to low APO concentrations but showed growth inhibition at concentrations ≥10 μM, while the growth of others was inhibited already at 5 μM APO. Generally, the impact of APO on growth was dose dependent (Figure 1A). In stark contrast to the overall weak impact of BOA, exposure to APO severely inhibited growth of 43% of isolates at concentrations of 50 μM: 77 out of 179 isolates grew to less than 50% of control levels in 50 μM APO (Figure 1A). All isolates in the collection that belonged to the families *Alcaligenaceae* (β-Proteobacteria) or *Pseudomonaceae* and *Moraxellaceae* (γ-Proteobacteria) were resistant to APO, while all *Flavobacteriaceae* (Bacteroidia) were APO sensitive. The remaining twenty families contained both sensitive and resistant isolates (Figure 1A, B).

From the overall phylogeny (Figure 1A), it appeared that there was some phylogenetic structuring of APO tolerance; within lower taxonomic ranks, however, this signal appeared to become less pronounced, indicating that APO tolerance might be a strain-rather than a species- or genus-specific trait in most families and orders (Figure 1B). To test this hypothesis, we calculated the strength of the correlation between the relative growth at 50 μM and the phylogenetic distance using Pagel’s λ and tested against the null hypothesis λ = 0 (likelihood ratio test, Supplemental Table 1; Pagel, 1999, Münkemüller et al., 2012). A value of λ close to 0 indicates no phylogenetic signal; conversely λ values closer to 1 indicate that the trait is correlated with the phylogenetic structure. For the overall collection, λ was 0.76 (p < 0.001). We next tested the strength of the phylogenetic signal within gram-positive bacteria (classes Actinomycetia and Bacilli), which returned a weaker but significant λ of 0.46 (p < 0.05). However, within both Actinomycetia and Bacilli, or within families belonging to those classes, this signal disappeared (λ < 0.1, p > 0.05). Within the phylum Proteobacteria (α-, β- and γ-proteobacteria), λ was high (0.81, p < 0.05). Within phyla, response correlated with phylogenetic structure only in β- and γ-proteobacteria (λ = 1, p < 0.05) but not in α-proteobacteria (λ < 0.1, p > 0.05). With the exception of the *Xanthomonadaceae* (λ = 1, p < 0.05), λ values within proteobacteria families were not significantly different from zero. Within the *Xanthomonadaceae*, where we observed a phylogenetic signal for APO response, four out of five sensitive isolates belonged to the genus *Lysobacter* (Supplemental Table 1, Supplemental Figure 1). These data corroborated our initial observation that APO tolerance is largely a strain-specific feature, and that diversity in this trait exists in the AtRSphere collection. While there was a phylogenetic signal in the dataset at broad phylogenetic and taxonomic levels, this signal often weakened or disappeared in lower taxonomic ranks (Figure 1A, B, Supplemental Table 1).

### Synthetic bacterial communities are structured by BX treatment

To study the effect of BX on communities, we designed four different SynComs based on the responses of the individual isolates: one that fully represented the isolate collection at the genus level by randomly choosing one isolate per genus (referred to hereafter as *random*) and three reduced-complexity SynComs (Supplemental Table 2). Of these, the *tolerant* SynCom only contained isolates (max. one per genus) that had grown to at least 50% of the control level in 50 μM APO. As can be seen in Figure 1A, several isolates that were part of *tolerant* were also included in *random*. The *sensitive* SynCom consisted only of isolates that had grown to less than 50% of control levels in 50 μM APO. Finally, the *mixed* SynCom contained both APO-sensitive and APO-tolerant isolates (Figure 1A). To understand the temporal and treatment-specific community dynamics, we grew each SynCom in three different environments: control (supplemented with DMSO, the solvent used in the other two treatments), 100 μM BOA, and 50 μM APO. We monitored SynCom composition in 24 h intervals over 4 d via 16S rRNA sequencing (see methods).

As all bacteria analyzed here had originally been isolated from complex plant root microbiomes, we first assessed to which degree they could reassemble into communities *in vitro* in the absence of a host. Constrained correspondence analysis (CCA) of Jaccard distances calculated from relative abundances revealed that *in vitro* SynComs assembled reproducibly (Figure 2A-D).

**Figure 2.**
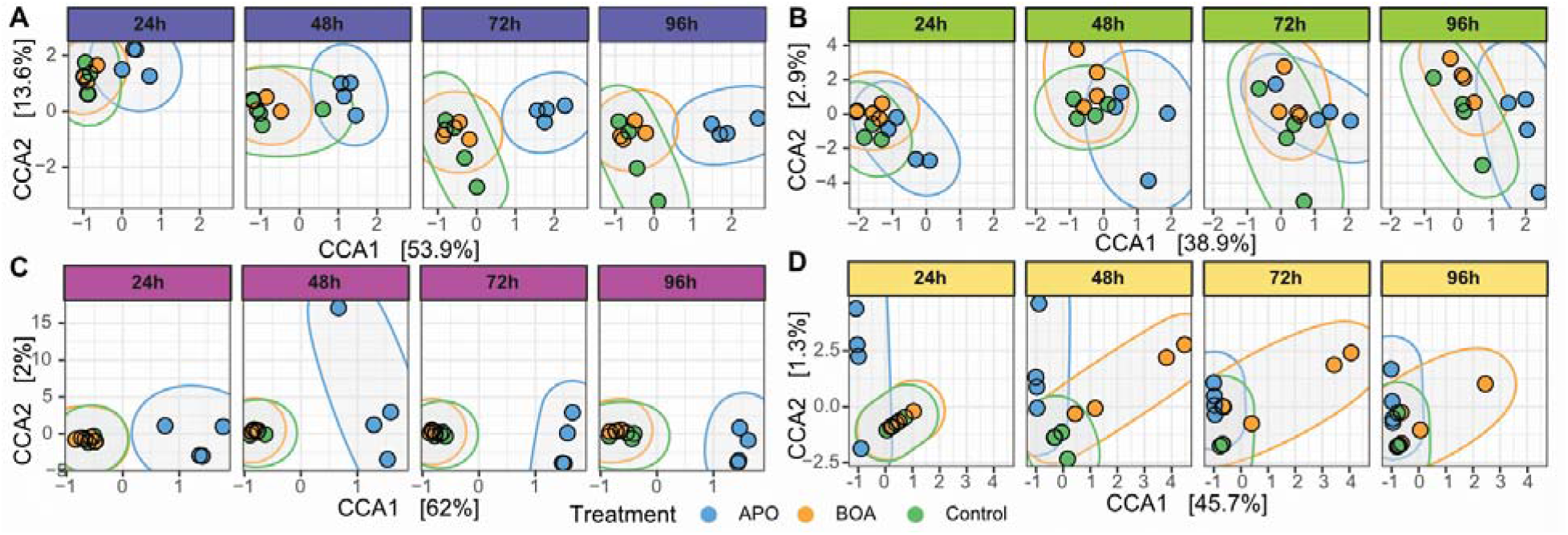
Community analysis. Constrained correspondence analysis (CCA, constrained by ‘Treatment’ and ‘Timepoint’) of the Jaccard distances computed on relative genus abundances for all samples. **A-D)** CCA per SynCom (**A**: random; **B**: tolerant; **C**: mixed; **D**: sensitive); faceted by time point (columns) and colored by treatment. **E)** Relationship between change in bacterial growth in the presence of BX compounds in pure culture (x-axis) plotted against the change in relative abundance when embedded in a community (y-axis) for APO (blue) and BOA (orange). Each dot shows the average change in relative abundance for one isolate across all time points.

We next assessed if communities containing only APO-sensitive isolates would be more strongly restructured by the addition of APO than one assembled only from APO-tolerant isolates. Generally, we observed treatment-specific changes in all SynComs. Overall, BOA-treated samples were similar to the control in all SynComs, with the exception of the *sensitive* one. APO-treated samples clustered separately from both control and BOA (Figure 2 A-D). The separation between APO and the remaining samples in the ordination was most obvious for the *mixed* and *random* and less striking for the *tolerant* SynCom; it decreased over time in the *sensitive* one (Figure 2 A-D).

We performed permutational multivariate analysis of variance (PERMANOVA) to see if we could confirm these observations. Treatment explained a significant proportion of variance in all SynComs: 40% in *random*, 22% in *tolerant*, 75% in *mixed*, and 41% in *sensitive*. This means that the proportion of between-sample variance explained by BX treatments was lowest in the community assembled from APO-tolerant isolates. Time point explained a significant proportion of variance in *random* and *tolerant* (31% and 33%, respectively) but not in *mixed* or *sensitive* (Supplemental Table 3). We then performed another PERMANOVA analysis on each SynCom, subset to a binary comparison between control and either APO or BOA treatment, to estimate the amount of variance explained by the individual treatments (Supplemental Table 4). This confirmed that most of the variance observed between treatments in each SynCom was driven by APO (Figure 2A-D). The amount of variance explained by APO treatment was lowest in the *tolerant* SynCom (22%), while APO treatment explained approx. 40% in *random* and *sensitive* and 70% in *mixed* (Supplemental Table 4). In other words, the SynCom composed of only the most APO-tolerant individual isolates showed the highest resilience to APO treatment.

Having confirmed that the different isolate mixtures reliably assembled into reproducible communities, we asked if within-sample species diversity (alpha-diversity) was affected by time or treatment by applying analysis of variance (ANOVA). In the *random, tolerant*, and *mixed* SynComs, APO (but not BOA) treatment significantly reduced the total number of taxa at one or more time points, while in the *sensitive* SynCom, the taxon number remained constant across treatments and time points (Supplemental Figure 2, Supplemental Tables 5 and 6).

Time alone, however, did not significantly influence the species number in any of the SynComs. The Shannon measure of alpha diversity, which in addition to the number of species also incorporates the evenness of species abundances, was significantly influenced by both time and treatment (Supplemental Figure 2, Supplemental Tables 5 and 6). While APO treatment significantly increased Shannon diversity of the *mixed* and *tolerant* communities, it decreased Shannon diversity of the *sensitive* one. In contrast, BOA significantly increased Shannon diversity of the *sensitive* SynCom after 48 h (Supplemental Tables 7 and 8).

### Isolates respond to APO in a community-specific fashion

Having substantiated that different SynComs display different resilience to BX treatment, we asked if the tolerance of an isolate to BX would predict how well this isolate could establish itself in a BX-treated community. Overall, there was no relationship between the relative growth of an individual isolate in treatment conditions and and treatment-dependent changes in relative abundance in a community (Figure 3A; adjusted R^2^ = 0.01 for APO and BOA, p > 0.05, Table S9). We also tested if there was a relationship between *absolute* growth in treatment and changes in relative abundance and found that absolute growth was not predictive of performance in a community either (adjusted R^2^ = 0.02 for APO, and R^2^ = −0.01 for BOA, p > 0.05, Table S9). In fact, some isolates that barely grew when cultured individually increased in relative abundance when grown in a treated community (Figure 3A, top left corner of the plot). After establishing that individual performance was a poor predictor of performance when embedded in a community, we asked if the ability of an isolate to establish itself in a BX-treated community instead depended on interactions with other community members. For this, we exploited the partial isolate overlap between the communities (Figure 1A). We assumed that if there were no community effects, an isolate would undergo the same treatment-related changes in every community, independent of social context. We therefore explored the treatment-specific changes in relative abundance for all isolates that were recovered in more than one community (Figure 3B). In total, 10 isolates that were included in more than one community significantly changed in relative abundance upon BX treatment (Figure 3B, FDR adjusted p < 0.05). Of these, seven isolates changed strongly (absolute log2FC > 1) in relative abundance in a treatment- and community-specific manner. For example, *Bacillus* Root239 did not change in relative abundance when the *tolerant* community was APO treated, but was strongly reduced under these conditions in the *mixed* community. Similarly, *Flavobacterium* Root420 was unaffected by APO treatment when embedded in the *mixed* community but was strongly reduced in relative abundance in the *random* community. *Acinetobacter* Root1280, *Bordetella* Root565 and *Noviherbaspirillum* Root189 underwent mild (absolute log2FC < 1) fold-changes in relative abundance when the communities they were embedded in were treated with APO. While for these three isolates the direction remained the same, the amplitude of the change was community-specific. This indicates that community-specific interactions between isolates can modulate the impact of chemical exposure on the individual isolate.

**Figure 3:**
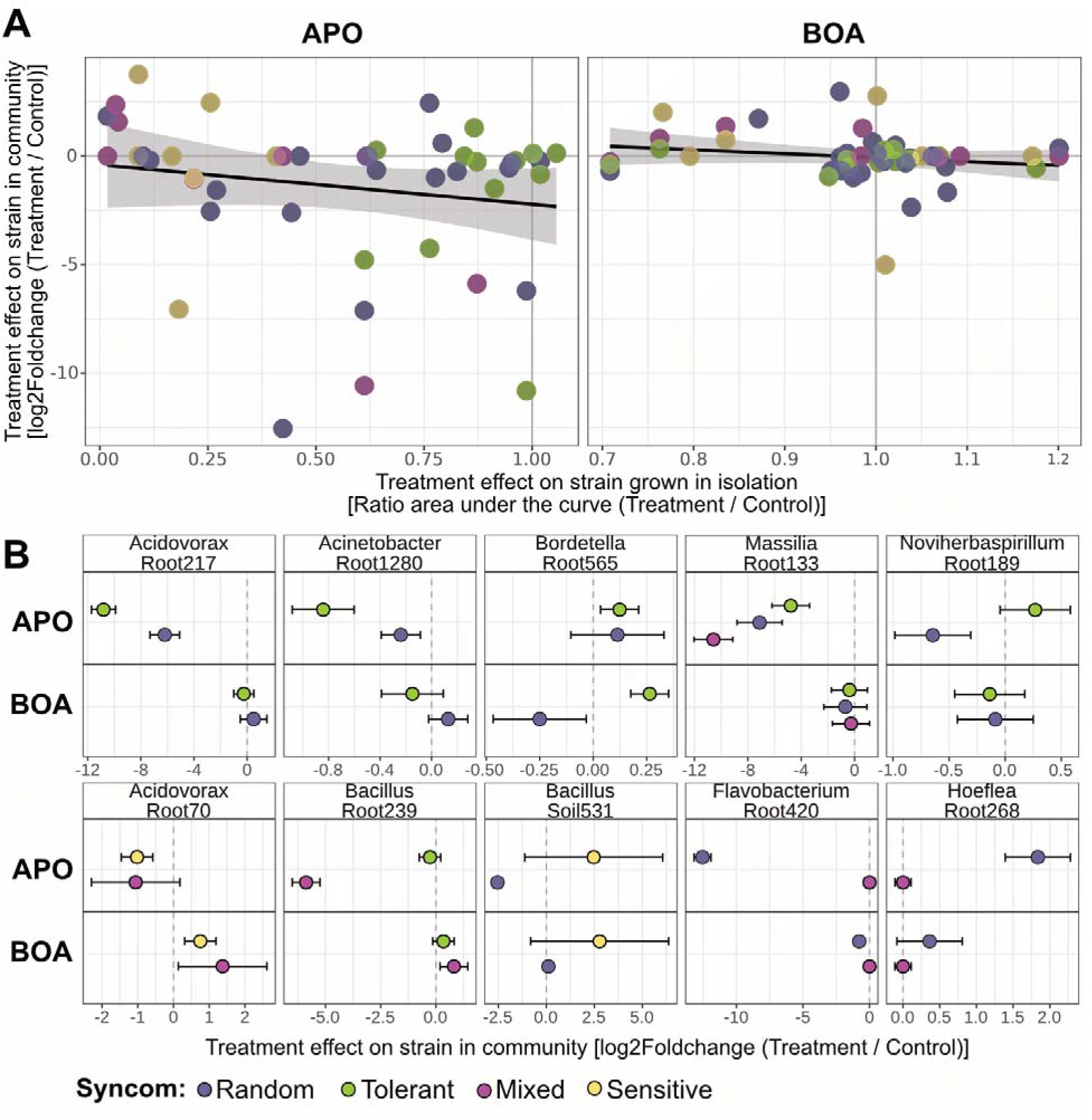
Comparison between isolate growth alone and in different communities. **A)** Dotplot of relative growth of an isolate in pure culture (ratio of area under the respective growth curves) on x plotted against log2(Foldchange) in relative abundance when embedded in a community (Treatment / Control) on y. The black line shows the regression line (y as the dependent variable, x as predictor) with 95% confidence intervals as grey shading. **B)** Dotplots of log2(Foldchange) in relative abundance by treatment for isolates found in multiple SynComs. Errors indicate s.e of the log2(Foldchange), colors indicate SynCom. Changes in relative abundance were calculated across all time points; a breakdown by time point is shown in Supplemental Figure 4.

However, it is important to consider not only the relative change per isolate but also the impact of this change on overall community composition. For example, an isolate dropping from 50% to 25% relative abundance (l2fc of −1), may be considered a more important change in the larger community context than an increase from 0.001% to 0.008% (l2fc of 3).

### Interactions between isolates are modulated by community context and chemical treatment

In the next approach, we wanted to further investigate which isolates interacted in a treatment- and community-specific manner. We focused our analysis on the *random* and *tolerant* communities, since these shared 5 highly abundant genera, allowing for a comparative approach (Figure 3B, Figure 4). To understand how SynCom-specific isolate interactions drove condition-specific compositional changes at the genus level, we plotted the relative abundance of highly abundant isolates (>1% average relative abundance) that belonged to genera shared between *random* and *tolerant* (Figure 4A).

**Figure 4:**
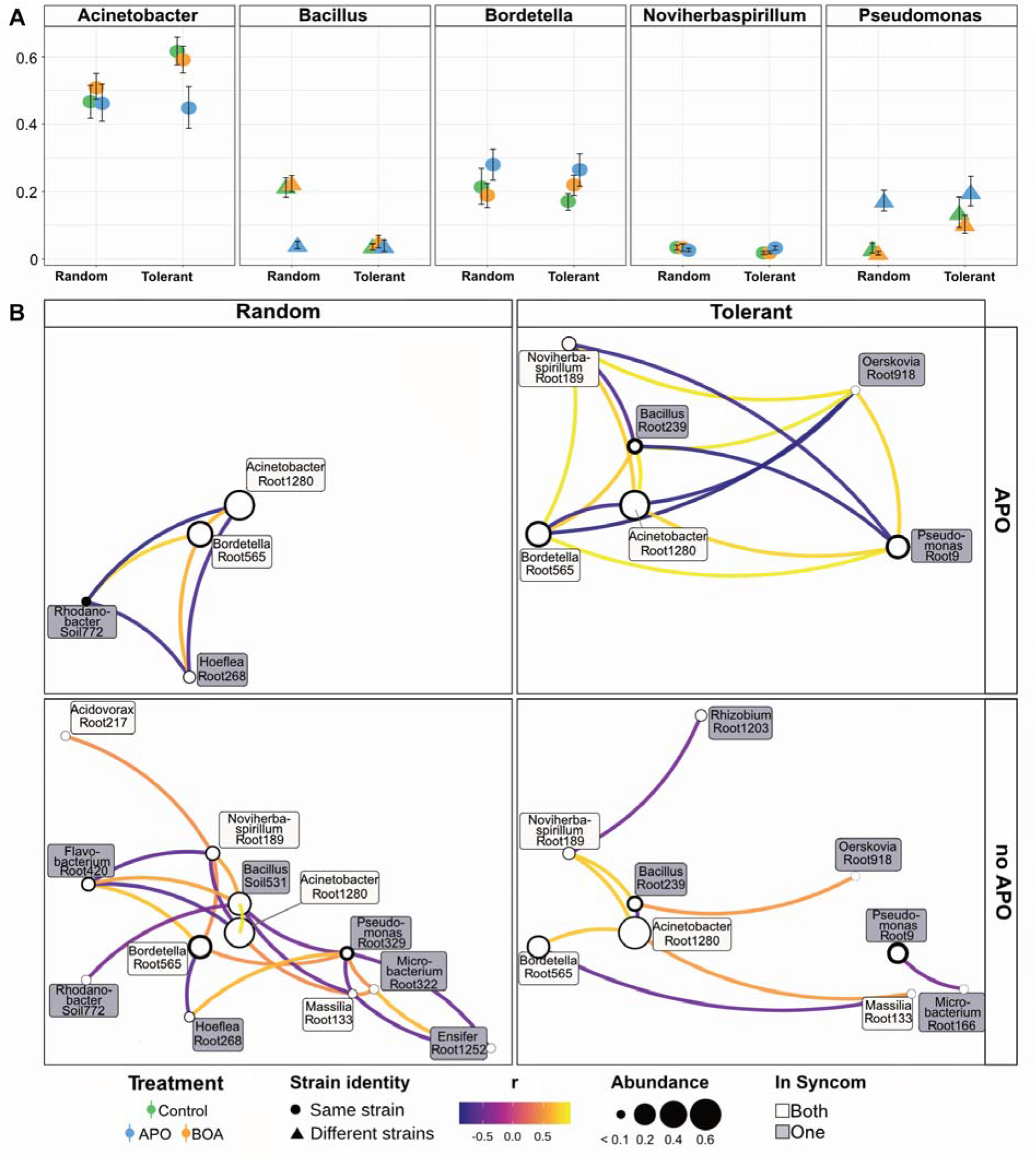
Differential isolate interactions in the *random* and *tolerant* SynComs are driven by isolate identity and chemical treatment. **A)** Relative abundance of Genera reaching more than 1% relative abundance and present in both SynComs. **B)** Partial Pearson’s correlations (FDR adjusted p < 0.1) of isolate abundances in the *tolerant* and *random* SynComs. Each node represents one isolate: the inner edge of the circle indicates mean-sd, the outer edge shows mean + sd of the relative abundance. Color of the node label indicates if an isolate was a member of both (white) or only one (grey) of the SynComs. Each edge indicates a significant correlation between two isolates; edge color indicates the correlation value, negative correlations have a blue hue, while positive correlations are orange to yellow.

In APO conditions, the relative abundances of these highly abundant genera were similar in both SynComs, irrespective of isolate identity (Figure 4A, blue shapes). However, when there was no chemical selection (i.e., in BOA or Control treatments), the communities diverged: the *Bacillus* isolate in *random* (Soil531) reached higher abundance than the one in *tolerant* (Root239). Conversely, the genera *Acinetobacter* (Root1280) and *Pseudomonas* (Root9) increased in abundance in *tolerant* compared with either the same strain (*Acinetobacter* Root1280) or a closely related counterpart (*Pseudomonas* Root329) in *random*. Both *Pseudomonas* isolates reached a lower relative abundance in control and BOA conditions than in APO treatment (Figure 3B, 4A), suggesting that *Pseudomonas* profited from APO irrespective of the community or strain identity, in line with the observation that *Pseudomonadaceae* in the collection were generally APO insensitive (Figure 1B). The effect of APO on *Bacillus* was strain- and community-specific: Root239 (*tolerant* SynCom) was lowly abundant in all SynCom / treatment combinations, while Soil531 (*random*) was low in APO-treated samples but reached 20% relative abundance in the samples that were not treated with APO. Correlation analysis using partial Pearson’s correlation coefficients revealed that the relative abundance of *Bacillus* Soil531 in the *random* SynCom was negatively correlated with the relative abundance of *Pseudomonas* Root329 in the absence of APO treatment (Figure 4B). This suggested that there was antagonism between these isolates: the high relative abundance of *Pseudomonas* Root329 in the APO-treated *random* community could be the result of a combination of its insensitivity to APO with an APO-dependent reduction of its antagonist *Bacillus* Soil531 (which was less abundant in APO). We also observed that both *Bacillus* isolates displayed community-specific chemical sensitivity (Figure 3A, Supplemental Figure 3): while *Bacillus* Root239 had low relative abundance in the *tolerant* community in all conditions, it reached >90% rel. abundance in BOA and control conditions when part of the *mixed* community (Figure 4A, Supplemental Figure 4). Similar to Soil531 in *random*, Root239 in *mixed* was strongly reduced in relative abundance upon APO treatment (Figure 3A), coinciding with increased relative abundances of *Hoeflea* Root268 and *Lysobacter* Root96 in the APO treated *mixed* community (Supplemental Figure 4), further highlighting how treatment-specific isolate interactions affect community composition.

## Discussion

### Diversity of BX sensitivity in culturable rhizosphere bacteria

In our screen of the *AtRSphere* collection for sensitivity to BX derivatives (Figure 1), we observed broad diversity of growth responses to BX exposure. While some clades were non-responsive to BOA and APO, we observed diverse responses in most clades. APO in particular exerted a strong negative influence on the growth of some isolates. These findings are in line with previous results showing the antibiotic activity of APO on anaerobic microorganisms (Atwal et al., 1992). In our study, BOA influenced bacterial growth only marginally compared to APO, similar to what has been described for their effects on fungi and plants. In general, this corroborates the previously suggested relevance of APO, which is more stable than BOA in soil, in BX-mediated plant-environment interactions (Kumar et al., 1993; Schandry and Becker, 2020; Schulz et al., 2013; Schütz et al., 2019; Venturelli et al., 2015; Voloshchuk et al., 2020). However, our data showed that bacterial response to APO was largely isolate-specific and not dictated by phylogenetic relationship (Figure 1B). This indicates that even within genera, natural genetic diversity can result in substantial differences in BX sensitivity. Indeed, previous studies reported functional overlap related to responses to plant-derived compounds between isolates from different clades, while not all isolates of the same clade shared these functionalities (Bai et al., 2015; Harbort et al., 2020). The broad phylogenetic spectrum of the *AtRSphere* collection prevented the identification of genetic components underlying bacterial response to BOA and APO. However, the strongly differential responses we observed among closely related isolates warrant a more comprehensive analysis of the genetic basis of the BX response in the respective clades in future experiments.

One major advantage of the *AtRSphere* collection is that it was isolated from a non-BX-producing host plant. The soil from which this collection was originally derived (Cologne soil) was collected from an agricultural site, and in absence of a complete record of the field’s history, we cannot exclude that there might have been an input of BX in the past. Our findings on the standing diversity of BX tolerance imply that this diversity is maintained in plant-associated bacteria, also in the absence of a BX-releasing host. Strain-specific tolerance or sensitivity to plant specialized metabolites may thus contribute to the colonization profiles observed for different plant species, where the composition is similar at higher taxonomic ranks but some isolates show host preference (Bulgarelli et al., 2013; Schneijderberg et al., 2020).

### Evaluation of bottom-up design strategy

Others have previously proposed that synthetic communities should be built with a focus on functional features instead of on phylogenetic coverage (Lemanceau et al., 2017; Vorholt et al., 2017). We employed a simple, function-guided strategy, compiling SynComs by binning individual isolates based on a specific phenotype and contrasting these binned assemblies to a randomly assembled SynCom. The decisive phenotype measured at the level of individual isolates and used for binning them into SynComs was growth upon exposure to APO. The phenotype measured at the community level, variance explained by treatment, was lowest (22%) in the SynCom assembled solely from APO-tolerant isolates, indicating that resistance of individual SynCom members leads to overall community resilience. While this approach worked well to create an overall resilient community, the amount of variance explained by treatment was similar in *random* and *sensitive* (~40%) and highest in the *mixed* SynCom (75%), indicating that mixing tolerant and sensitive isolates does not necessarily result in a community with intermediate resilience.

### BX dependent community alterations *in vitro* mimic those described in soil

Our work complements recent research efforts to understand the effect of benzoxazinoids on plant-associated microbiota in soil (Cadot et al., 2020; Cotton et al., 2019; Hu et al., 2018; Kudjordjie et al., 2019; Neal et al., 2012). Ultimately, for *in vitro* SynCom systems to be a useful model for plant microbiota research, they should behave similarly to microbiota in soil. BX constitute a prime system to assess this, as the influence of BX release on the root and rhizosphere microbiome has been investigated via mutant studies in different soils (Cadot et al., 2020; Cotton et al., 2019; Hu et al., 2018; Kudjordjie et al., 2019). Among the bacterial taxa that were found in different studies, Flavobacteriales were observed as depleted in multiple studies. Hu et al (Hu et al., 2018) found *Flavobacterium* depleted on the roots of BX-exuding plants and the surrounding soils when comparing wildtype and *bx1* mutant plants. Extending these observations, Cadot et al (Cadot et al., 2020) reported that *Flavobacteriaceae* were consistently depleted on roots of BX-producing plants, independent of soil type and location. Similarly, members of the Flavobacteriales family were more abundant on the roots of maize BX-synthesis mutants than on wildtype in another, independent study (Cotton et al., 2019). In all of the afore-mentioned studies, *Comamonadaceae* were also reported to be depleted on roots of BX-releasing plants when compared to BX-deficient mutants (Cadot et al., 2020; Cotton et al., 2019; Hu et al., 2018; Kudjordjie et al., 2019).

Our data show that the growth of *Flavobacterium* isolates was only slightly inhibited in growth across APO levels, with some isolates displaying mild sensitivity to BOA (Figure 1). However, *Flavobacterium* was significantly depleted from the *random* and *sensitive* SynComs once the community was exposed to APO (Supplemental Figure 4). Similarly, *Acidovorax* (*Comamonadaceae*) was found to be reduced in the APO treatment in all SynComs. Thus, even though *in vitro* cultivation strongly differs from soils in physical and chemical parameters and in overall complexity, our study suggests that plant-exudate-mediated effects on soil communities can be recapitulated *in vitro*, highlighting the applicability of such reduced-complexity systems for future studies. In addition, such simplified systems may aid in the identification of microbial isolates that act as community keystones, or hubs, under specific chemical challenges in a short experimental time frame (Carlström et al., 2019).

### Chemical stability of isolate networks

One might expect that in an *in vitro* system containing a mixture of isolates, one isolate outgrows the others and reaches very high abundance. Under that assumption, there would be no interactions between the individual isolates and growth rate would be a strong predictor of final abundance. However, we found that the growth of an isolate by itself was only poorly correlated with its ability to establish itself as a SynCom member *in vitro* or with its relative abundance in a community (Figure 3A). In fact, several isolates that grew poorly when cultured alone reached high relative abundance when grown in communities, possibly indicating facilitation by other members or other types of positive interactions (Sanchez-Gorostiaga, et al., 2019)

We found that some isolates that reached only low relative abundances in control communities were able to establish themselves much better when the community was treated with APO (Figure 4, Supplemental Figure 4). We propose that the positive effects of APO on certain community members is not due to a direct positive effect of APO, but instead due to a negative effect of APO on a competitor or antagonist in the same community. Interestingly, we have observed two instances where the APO-dependent reduction of a *Bacillus* isolate coincided with improved growth of other isolates (Figure 4, Supplemental Figure 4). *Bacillus* species are generally competitive, and often produce a range of compounds that inhibit the growth of other bacteria (Shafi et al., 2017) lending further support to the hypothesis that a negative impact of APO on *Bacillus* isolates contributes to the APO-specific changes in community composition. *Bacillus* members thus seem to shape the synthetic communities in a manner opposite to a “keystone”, whose presence typically facilitates the growth of other community members. This phenomenon has been previously observed for *Bacillus* species and probably contributes to the *Bacillus*-mediated biocontrol against certain plant-pathogens observed in natural soils (Lee et al., 2021).

## Methods

### Isolate profiling

Individual isolates were pre-cultured in ½ strength TSB medium in 96-Well 2 ml deep-well plates (Semadeni) covered with a Breathe-Easy (Diversified Biotech) membrane until stationary phase for 6 d at 28°C and 180 rpm. Cultures were then resuspended by pipetting up and down 10 times and inoculated into 96-well flat-bottom, transparent cell culture plates (Nunc), containing either solvent control (DMSO), 1 μM APO, 5 μM APO, 10 μM APO, 50 μM APO, 10 μM BOA, 50 μM BOA or 100 μM BOA in ½-strength TSB. Different media were prepared by mixing a stock containing the highest concentration of the respective chemical, with medium containing the solvent at the same concentration (DMSO; 1:20.000). Pipetting was carried out using an Agilent Bravo pipetting robot equipped with a 96-tip head. Plates were stacked without lids, with one empty plate at the top and the bottom of the stack and placed into a BioTek plate stacker connected to a BioTek Synergy2 plate reader. Plates were shaken for 120 s before reading, and absorption (600 nm) was measured in each well. Plates were continuously shaken, measured, and stacked over the course of at least 48 h per replicate. For each replicate, the relative growth per isolate and condition was calculated by dividing the area under the growth curve (x = time in minutes; y = increase in OD600) in the respective condition by the area under the growth curve of the control condition. The experiment was carried out in eight replicates per condition.

### Synthetic bacterial communities

#### Design

To investigate to which degree the overall resilience of a bacterial community to chemical changes is determined by the properties of its individual members, we followed two separate design strategies. In the first strategy, we aimed at creating a diverse community spanning all genera of the isolate collection. The *random* community was created by picking one random isolate per genus.

In the second approach, we built communities based on the growth profiles of the individual isolates. These communities span a limited number of genera, but the member growth profile is more strictly defined than in the “full” community. One of the reduced representation communities was created from isolates that grew to 75% to 100% of control conditions in 50μM APO (*tolerant*). The other reduced community was assembled from isolates that grew to 0 to 25% of control in 50μM APO (*sensitive*). If there was more than one isolate per genus that fulfilled the condition, we selected one at random. In addition, we compiled one limited representation SynCom containing a mix of sensitive and tolerant isolates (*mixed*).

### Cultivation and sequencing

Individual isolates were pre-cultured in ½-strength TSB medium in 96-Well 2ml deep-well plates covered with a Breathe-Easy membrane (Diversified Biotech) with shaking at 28°C for 6 days until stationary phase, as described previously (Bai et al., 2015). Individual cultures were resuspended by pipetting up and down 10 times, and then 100 μL of each culture were mixed with 10 mL ½-strength TSB medium containing solvent at the same concentration as the treatment cultures (1:20.000 DMSO). 100 μL of the mixed culture were used to inoculate 20 mL cultures containing either a solvent control (DMSO), 100 μM BOA, or 50 μM APO in 100 mL Erlenmayer flasks. Flasks were cultured at 28°C, 180 rpm; 1 ml culture was sampled every 24 h. Sampled cultures were centrifuged at 12.000 rpm, the supernatant was discarded and the pellet was stored at −70 °C until further use. DNA was extracted using the DNA PowerSoil kit (Qiagen) following the standard protocol. 16S amplicons were generated according to the Illumina 16S protocol, indexed using Nextera XT dual indexing (Illumina), and pooled per time point for sequencing on a MiSeq (Paired end 300bp, Illumina).

### Data analysis

RStudio with R version 4.0.2 was used for data analysis. The phylogenetic tree in Figure 1 was computed from concatenated 16SrRNA, gyrB, gyrB, rpoB, recA, atpD, dnaJ and thrC DNA sequences. These sequences were extracted from the fasta files containing nucleotide ORFs (obtained from https://at-sphere.com/) using tBLASTn of the respective *E. coli* protein sequence. The concatenated sequences were aligned using the DECIPHER R package (Wright, 2016) and a maximum likelihood phylogeny with the general time reversible model was fitted using the phangorn package (Schliep et al., 2017; Schliep 2011). Pagel’s λ and the likelihood ratio test results were obtained using the phylosig function from phytools (Revell, 2012). For the analysis of 16S amplicon data, reads were processed and ASVs were inferred using the dada2 pipeline (Callahan et al., 2016). Taxonomic assignment of ASVs was done using SynCom specific 16SrRNA databases and agglomerated at the genus level. ASVs that could not be assigned were discarded. Partial Pearson’s correlation coefficients were calculated and tested for isolates with >100 reads per sample using the correlate R package. Scripts used for data analysis are available at https://github.com/nschan/BxSynComMs. The following R packages were used with R version 4.0.2: Tidyverse (Wickham et al., 2019), Broom (Robinson et al., 2019), dada2 (Callahan et al., 2016), DECIPHER (Wright, 2016), DESeq2 (Love et al., 2014), emmeans (Lenth et al., 2019), ggforce (Pedersen, 2020a), ggthemes (Arnold, 2019), ggtree (Yu, 2020; Yu et al., 2018, 2017), gt (Iannone et al., 2019), igraph (Csardi and Nepusz, 2006), MESS (Ekstrøm, 2016), multcomp (Hothorn et al., 2008), patchwork (Pedersen, 2020b), phangorn (Schliep et al., 2017; Schliep, 2011), phyloseq (McMurdie and Holmes, 2013), phytools (Revell, 2012), vegan (Oksanen et al., 2019), wesanderson (Ram and Wickham, 2018) in combination with some custom functions.

### Data availability

Sequencing reads have been deposited to the European Nucleotide Archive (ENA; www.ebi.ac.uk) under the accession no. PRJEB42439.

## Acknowledgements

The authors gratefully acknowledge support from the Vienna BioCenter Core Facility (VBCF) Molecular Biology Services, especially Dr. Robert Heinen, the VBCF Media Kitchen, the VBCF Sequencing facility and the VBCF High Performance Computing facility. The authors also gratefully acknowledge the Leibniz Supercomputing Centre (LRZ) for funding this project by providing computing time on its Linux-Cluster and Rstudio Server.

The authors thank Patrick Hüther, Talia Karasov and Eva Knoch for helpful discussion and constructive feedback on the manuscript, and Iacopo Gentile and Karina Weiser-Lobao for conducting preliminary experiments and for technical support. The authors also thank Stijn Staepen for help with the *AtRSphere* collection and Paul Schulze-Lefert for helpful discussion.

This work was funded by the Austrian Academy of Sciences (ÖAW) and the European Research Council (ERC) under the European Union’s Horizon 2020 research and innovation programme (Grant agreement No. 716823, “FEAR-SAP”).

## Supplemental Material

The following supplemental material is contained in the file

Schandry_et_al_supplemental_material.pdf

## Supplemental Tables

**Supplemental Table 1. Phylogenetic signal**

**Supplemental Table 2. Syncom strains**

**Supplemental Table 3. PERMANOVA**

**Supplemental Table 4 PERMANOVA on treatment subsets**

**Supplemental Table 5. ANOVA: observed taxa**

**Supplemental Table 6. Observed diversity means and pairwise comparisons**

**Supplemental Table 7. ANOVA: Shannon Diversity**

**Supplemental Table 8. Shannon diversity means and pairwise comparisons**

**Supplemental Table 9. Correlation between change in relative abundance and change in growth**

## Supplemental figures

**Supplemental Figure 1. Bx sensitivity of Xanthomonadales**

**Supplemental Figure 2. Observed alpha diversity**

**Supplemental Figure 3. Shannon diversity indices of the synthetic communities**

**Supplemental Figure 4. Changes in relative abundance by community**

## Literature

Anzai K, Isono K, Okuma K, Suzuki S. 1960. The new antibiotics, questiomycins A and B. J Antibiot 13:125–132.

Arnold JB. 2019. ggthemes: Extra Themes, Scales and Geoms for “ggplot2.”

Atwal AS, Teather RM, Liss SN, Collins FW. 1992. Antimicrobial activity of 2-aminophenoxazin-3-one under anaerobic conditions. Can J Microbiol 38:1084–1088.

Bai Y, Müller DB, Srinivas G, Garrido-Oter R, Potthoff E, Rott M, Dombrowski N, Münch PC, Spaepen S, Remus-Emsermann M, Hüttel B, McHardy AC, Vorholt JA, Schulze-Lefert P. 2015. Functional overlap of the Arabidopsis leaf and root microbiota. Nature 1–19.

Banerjee S, Schlaeppi K, van der Heijden MGA. 2018. Keystone taxa as drivers of microbiome structure and functioning. Nat Rev Microbiol 16:567–576.

Belz RG, Hurle K. 2005. Differential exudation of two benzoxazinoids--one of the determining factors for seedling allelopathy of Triticeae species. J Agric Food Chem 53:250–261.

Bodenhausen N, Bortfeld-Miller M, Ackermann M, Vorholt JA. 2014. A synthetic community approach reveals plant genotypes affecting the phyllosphere microbiota. PLoS Genet 10:e1004283.

Bulgarelli D, Rott M, Schlaeppi K, Ver Loren van Themaat E, Ahmadinejad N, Assenza F, Rauf P, Huettel B, Reinhardt R, Schmelzer E, Peplies J, Gloeckner FO, Amann R, Eickhorst T, Schulze-Lefert P. 2012. Revealing structure and assembly cues for Arabidopsis root-inhabiting bacterial microbiota. Nature 488:91–95.

Bulgarelli D, Schlaeppi K, Spaepen S, van Themaat EVL, Schulze-Lefert P. 2013. Structure and Functions of the Bacterial Microbiota of Plants. Annu Rev Plant Biol 64:807–838.

Cadot S, Guan H, Bigalke M, Walser J-C, Jander G, Erb M, van der Heijden M, Schlaeppi K. 2020. Specific and conserved patterns of microbiota-structuring by maize benzoxazinoids in the field. bioRxiv. doi:10.1101/2020.05.03.075135

Callahan BJ, McMurdie PJ, Rosen MJ, Han AW, Johnson AJA, Holmes SP. 2016. DADA2: High-resolution sample inference from Illumina amplicon data. Nat Methods 13:581–583.

Callahan BJ, Sankaran K, Fukuyama JA, McMurdie PJ, Holmes SP. 2016. Bioconductor Workflow for Microbiome Data Analysis: from raw reads to community analyses. F1000Res 5:1492.

Carlström CI, Field CM, Bortfeld-Miller M, Müller B, Sunagawa S, Vorholt JA. 2019. Synthetic microbiota reveal priority effects and keystone strains in the Arabidopsis phyllosphere. Nat Ecol Evol 3:1445–1454.

Cheng C, Othman EM, Fekete A, Krischke M, Stopper H, Edrada-Ebel R, Mueller MJ, Hentschel U, Abdelmohsen UR. 2016. Strepoxazine A, a new cytotoxic phenoxazin from the marine sponge-derived bacterium Streptomyces sp. SBT345. Tetrahedron Lett 57:4196–4199.

Cotton TEA, Pétriacq P, Cameron DD, Meselmani MA, Schwarzenbacher R, Rolfe SA, Ton J. 2019. Metabolic regulation of the maize rhizobiome by benzoxazinoids. ISME J. doi:10.1038/s41396-019-0375-2

Csardi G, Nepusz T. 2006. The igraph software package for complex network research. InterJournal.

De Roy K, Marzorati M, Van den Abbeele P, Van de Wiele T, Boon N. 2014. Synthetic microbial ecosystems: an exciting tool to understand and apply microbial communities. Environ Microbiol 16:1472–1481.

Dessaux Y, Grandclément C, Faure D. 2016. Engineering the Rhizosphere. Trends Plant Sci 21:266–278.

Ekstrøm C. 2016. MESS: Miscellaneous Esoteric Statistical Scripts.

Finkel OM, Salas-González I, Castrillo G, Conway JM, Law TF, Teixeira PJPL, Wilson ED, Fitzpatrick CR, Jones CD, Dangl JL. 2020. A single bacterial genus maintains root growth in a complex microbiome. Nature. doi:10.1038/s41586-020-2778-7

Fitzpatrick CR, Salas-González I, Conway JM, Finkel OM, Gilbert S, Russ D, Teixeira PJPL, Dangl JL. 2020. The Plant Microbiome: From Ecology to Reductionism and Beyond. Annu Rev Microbiol. doi:10.1146/annurev-micro-022620-014327

Frey M, Chomet P, Glawischnig E, Stettner C, Grün S, Winklmair A, Eisenreich W, Bacher A, Meeley RB, Briggs SP, Simcox K, Gierl A. 1997. Analysis of a chemical plant defense mechanism in grasses. Science 277:696–699.

Handrick V, Robert CAM, Ahern KR, Zhou S, Machado RAR, Maag D, Glauser G, Fernandez-Penny FE, Chandran JN, Rodgers-Melnik E, Schneider B, Buckler ES, Boland W, Gershenzon J, Jander G, Erb M, Köllner TG. 2016. Biosynthesis of 8-O-Methylated Benzoxazinoid Defense Compounds in Maize. Plant Cell 28:1682–1700.

Harbort CJ, Hashimoto M, Inoue H, Niu Y, Guan R, Rombolà AD, Kopriva S, Mathias J E E, Sattely ES, Garrido-Oter R, Schulze-Lefert P. 2020. Root-Secreted Coumarins and the Microbiota Interact to Improve Iron Nutrition in Arabidopsis. Cell Host Microbe 0. doi:10.1016/j.chom.2020.09.006

Herrera Paredes S, Gao T, Law TF, Finkel OM, Mucyn T, Teixeira PJPL, Salas González I, Feltcher ME, Powers MJ, Shank EA, Jones CD, Jojic V, Dangl JL, Castrillo G. 2018. Design of synthetic bacterial communities for predictable plant phenotypes. PLoS Biol 16:e2003962.

Hothorn T, Bretz F, Westfall P. 2008. Simultaneous inference in general parametric models. Biom J 50:346–363.

Hu L, Mateo P, Ye M, Zhang X, Berset JD, Handrick V, Radisch D, Grabe V, Köllner TG, Gershenzon J, Robert CAM, Erb M. 2018. Plant iron acquisition strategy exploited by an insect herbivore. Science 361:694–697.

Hu L, Robert CAM, Cadot S, Zhang X, Ye M, Li B, Manzo D, Chervet N, Steinger T, van der Heijden MGA, Schlaeppi K, Erb M. 2018. Root exudate metabolites drive plant-soil feedbacks on growth and defense by shaping the rhizosphere microbiota. Nat Commun 9:2738.

Iannone R, Cheng J, Schloerke B. 2019. gt: Easily Create Presentation-Ready Display Tables. R package version 0.1.0.

Imai S, Shimazu A, Furihata K, Furihata K, Hayakawa Y, Seto H. 1990. Isolation and structure of a new phenoxazine antibiotic, exfoliazone, produced by Streptomyces exfoliatus. J Antibiot 43:1606–1607.

Jacoby RP, Koprivova A, Kopriva S. 2020. Pinpointing secondary metabolites that shape the composition and function of the plant microbiome. J Exp Bot. doi:10.1093/jxb/eraa424

Ke J, Wang B, Yoshikuni Y. 2020. Microbiome Engineering: Synthetic Biology of Plant-Associated Microbiomes in Sustainable Agriculture. Trends Biotechnol. doi:10.1016/j.tibtech.2020.07.008

Koprivova A, Schuck S, Jacoby RP, Klinkhammer I, Welter B, Leson L, Martyn A, Nauen J, Grabenhorst N, Mandelkow JF, Zuccaro A, Zeier J, Kopriva S. 2019. Root-specific camalexin biosynthesis controls the plant growth-promoting effects of multiple bacterial strains. Proc Natl Acad Sci U S A. doi:10.1073/pnas.1818604116

Kudjordjie EN, Sapkota R, Steffensen SK, Fomsgaard IS, Nicolaisen M. 2019. Maize synthesized benzoxazinoids affect the host associated microbiome. Microbiome 7:59.

Kumar P, Gagliardo RW, Chilton WS. 1993. Soil transformation of wheat and corn metabolites mboa and DIM2BOA into aminophenoxazinones. J Chem Ecol 19:2453–2461.

Lebeis SL, Paredes SH, Lundberg DS, Breakfield N, Gehring J, McDonald M, Malfatti S, Glavina del Rio T, Jones CD, Tringe SG, Dangl JL. 2015. Salicylic acid modulates colonization of the root microbiome by specific bacterial taxa. Science 349:860–864.

Lee, S.-M., Kong, H.G., Song, G.C., Ryu, C.-M. 2021) Disruption of Firmicutes and Actinobacteria abundance in tomato rhizosphere causes the incidence of bacterial wilt disease. ISME J. 15: 330–347

Lemanceau P, Blouin M, Muller D, Moënne-Loccoz Y. 2017. Let the Core Microbiota Be Functional. Trends Plant Sci 22:583–595.

Lenth R, Singmann H, Love J, Buerkner P, Herve M. 2019. emmeans: Estimated Marginal Means, aka Least-Squares Means (Version 1.3.4).

Li L, Abou-Samra E, Ning Z, Zhang X, Mayne J, Wang J, Cheng K, Walker K, Stintzi A, Figeys D. 2019. An in vitro model maintaining taxon-specific functional activities of the gut microbiome. Nat Commun 10:4146.

Liu Y-X, Qin Y, Bai Y. 2019. Reductionist synthetic community approaches in root microbiome research. Curr Opin Microbiol 49:97–102.

Love MI, Huber W, Anders S. 2014. Moderated estimation of fold change and dispersion for RNA-seq data with DESeq2. Genome Biol 15:550.

Macías FA, Oliveros-Bastidas A, Marín D, Castellano D, Simonet AM, Molinillo JMG. 2005. Degradation studies on benzoxazinoids. Soil degradation dynamics of (2R)-2-O-beta-D-glucopyranosyl-4-hydroxy-(2H)-1,4-benzoxazin-3(4H)-one (DIBOA-Glc) and its degradation products, phytotoxic allelochemicals from Gramineae. J Agric Food Chem 53:554–561.

Macías FA, Oliveros-Bastidas A, Marín D, Castellano D, Simonet AM, Molinillo JMG. 2004. Degradation Studies on Benzoxazinoids. Soil Degradation Dynamics of 2,4-Dihydroxy-7-methoxy-(2 H)-1,4-benzoxazin-3(4 H)-one (DIMBOA) and Its Degradation Products, Phytotoxic Allelochemicals from Gramineae. J Agric Food Chem 52:6402–6413.

Maier L, Pruteanu M, Kuhn M, Zeller G, Telzerow A, Anderson EE, Brochado AR, Fernandez KC, Dose H, Mori H, Patil KR, Bork P, Typas A. 2018. Extensive impact of non-antibiotic drugs on human gut bacteria. Nature 555:623–628.

McMurdie PJ, Holmes S. 2013. phyloseq: an R package for reproducible interactive analysis and graphics of microbiome census data. PLoS One 8:e61217.

Münkemüller, T., Lavergne, S., Bzeznik, B., Dray, S., Jombart, T., Schiffers, K., and Thuiller, W. 2012. How to measure and test phylogenetic signal: How to measure and test phylogenetic signal. Methods Ecol. Evol. 3: 743–756.

Neal AL, Ahmad S, Gordon-Weeks R, Ton J. 2012. Benzoxazinoids in root exudates of maize attract Pseudomonas putida to the rhizosphere. PLoS One 7:e35498.

Oksanen J, Blanchet FG, Friendly M, Kindt R, Legendre P, McGlinn D, Minchin PR, O’Hara RB, Simpson GL, Solymos P, Stevens MHH, Szoecs E, Wagner H. 2019. vegan: Community Ecology Package.

Pagel, M. 1999. Inferring the historical patterns of biological evolution. Nature 401: 877–884.

Pedersen TL. 2020a. ggforce: Accelerating “ggplot2.”

Pedersen TL. 2020b. patchwork: The Composer of Plots.

Ram K, Wickham H. 2018. wesanderson: A Wes Anderson Palette Generator.

Revell, L. J. 2012. phytools: An R package for phylogenetic comparative biology (and other things). Methods Ecol. Evol. 3 217–223.

Robert CAM, Veyrat N, Glauser G, Marti G, Doyen GR, Villard N, Gaillard MDP, Köllner TG, Giron D, Body M, Babst BA, Ferrieri RA, Turlings TCJ, Erb M. 2011. A specialist root herbivore exploits defensive metabolites to locate nutritious tissues. Ecol Lett 14:55–64.

Robinson D, Hayes A, Couch S. 2019. Broom: Convert statistical objects into tidy tibbles.

Sanchez-Gorostiaga, A., Bajić, D., Osborne, M.L., Poyatos, J.F., Sanchez, A. 2019. High-order interactions distort the functional landscape of microbial consortia. PLoS Biol. 17:e3000550.

Schandry N, Becker C. 2020. Allelopathic Plants: Models for Studying Plant–Interkingdom Interactions. Trends Plant Sci 25:176–185.

Schliep, Klaus, Potts, J. A, Morrison, A. D, Grimm, W. G. 2017. Intertwining phylogenetic trees and networks. Methods in Ecology and Evolution. doi:10.1111/2041-210X.12760

Schliep KP. 2011. phangorn: phylogenetic analysis in R. Bioinformatics. doi:10.1093/bioinformatics/btq706

Schneijderberg M, Cheng X, Franken C, de Hollander M, van Velzen R, Schmitz L, Heinen R, Geurts R, van der Putten WH, Bezemer TM, Bisseling T. 2020. Quantitative comparison between the rhizosphere effect of Arabidopsis thaliana and co-occurring plant species with a longer life history. ISME J. doi:10.1038/s41396-020-0695-2

Schulz M, Marocco A, Tabaglio V, Macias FA, Molinillo JMG. 2013. Benzoxazinoids in rye allelopathy - from discovery to application in sustainable weed control and organic farming. J Chem Ecol 39:154–174.

Schütz V, Bigler L, Girel S, Laschke L, Sicker D, Schulz M. 2019. Conversions of Benzoxazinoids and Downstream Metabolites by Soil Microorganisms. Frontiers in Ecology and Evolution 7:238.

Shafi, J., Tian, H., and Ji, M. 2017. Bacillus species as versatile weapons for plant pathogens: a review. Biotechnol. Biotechnol. Equip. 31: 446–459.

Stringlis IA, Yu K, Feussner K, de Jonge R, Van Bentum S, Van Verk MC, Berendsen RL, Bakker PAHM, Feussner I, Pieterse CMJ. 2018. MYB72-dependent coumarin exudation shapes root microbiome assembly to promote plant health. Proc Natl Acad Sci U S A. doi:10.1073/pnas.1722335115

Trivedi P, Leach JE, Tringe SG, Sa T, Singh BK. 2020. Plant-microbiome interactions: from community assembly to plant health. Nat Rev Microbiol. doi:10.1038/s41579-020-0412-1

Venturelli S, Belz RG, Kamper A, Berger A, von Horn K, Wegner A, Bocker A, Zabulon G, Langenecker T, Kohlbacher O, Barneche F, Weigel D, Lauer UM, Bitzer M, Becker C. 2015. Plants Release Precursors of Histone Deacetylase Inhibitors to Suppress Growth of Competitors. Plant Cell 27:3175–3189.

Venturelli S, Petersen S, Langenecker T, Weigel D, Lauer UM, Becker C. 2016. Allelochemicals of the phenoxazinone class act at physiologically relevant concentrations. Plant Signal Behav 11:e1176818.

Voges MJEEE, Bai Y, Schulze-Lefert P, Sattely ES. 2019. Plant-derived coumarins shape the composition of an Arabidopsis synthetic root microbiome. Proc Natl Acad Sci U S A 116:12558–12565.

Voloshchuk N, Schütz V, Laschke L, Gryganskyi AP, Schulz M. 2020. The Trichoderma viride F-00612 consortium tolerates 2-amino-3H-phenoxazin-3-one and degrades nitrated benzo[d]oxazol-2(3H)-one. Chemoecology. doi:10.1007/s00049-020-00300-w

Vorholt JA, Vogel C, Carlström CI, Müller DB. 2017. Establishing Causality: Opportunities of Synthetic Communities for Plant Microbiome Research. Cell Host Microbe 22:142–155.

Vrancken G, Gregory AC, Huys GRB, Faust K, Raes J. 2019. Synthetic ecology of the human gut microbiota. Nat Rev Microbiol 17:754–763.

Wickham H, Averick M, Bryan J, Chang W, McGowan L, François R, Grolemund G, Hayes A, Henry L, Hester J, Kuhn M, Pedersen T, Miller E, Bache S, Müller K, Ooms J, Robinson D, Seidel D, Spinu V, Takahashi K, Vaughan D, Wilke C, Woo K, Yutani H. 2019. Welcome to the Tidyverse. JOSS 4:1686.

Wright ES. 2016. Using DECIPHER v2. 0 to analyze big biological sequence data in R. R J 8.

Yu G. 2020. Using ggtree to Visualize Data on Tree-Like Structures. Current Protocols in Bioinformatics. doi:10.1002/cpbi.96

Yu G, Lam TT-Y, Zhu H, Guan Y. 2018. Two methods for mapping and visualizing associated data on phylogeny using ggtree. Molecular Biology and Evolution. doi:10.1093/molbev/msy194

Yu G, Smith D, Zhu H, Guan Y, Lam TT-Y. 2017. ggtree: an R package for visualization and annotation of phylogenetic trees with their covariates and other associated data. Methods in Ecology and Evolution. doi:10.1111/2041-210X.12628

