## Supplemental Material for "Plant-derived benzoxazinoids act as antibiotics and shape bacterial communities"

### Supplemental Tables

#### Supplemental Table 1. Phylogenetic signal lambda

| Class | Order | Family | lambda | p.value |
|---|---|---|---|---|
| All - All | | | | |
| All | All | All | 0.763 | 0.0004 * * |
| Gram Negative - Proteobacteria | | | | |
| All | All | All | 0.817 | 0.0017 * * |
| Alpha | All | All | 0 | 0.9998 |
| Alpha | Rhizobiales | All | 0 | 1.0000 |
| Alpha | Rhizobiales | Phyllobacteriacae | 0 | 1.0000 |
| Alpha | Rhizobiales | Rhizobiacaea | 0.275 | 0.8837 |
| Beta | All | All | 1.034 | 0.0036 * * |
| Beta | Burkholderiales | All | 0.147 | 0.6171 |
| Beta | Burkholderiales | Comamonadaceae | 0 | 1.0000 |
| Gamma | All | All | 1 | 0.0000 * * |
| Gamma | Xanthomonadales | All | 1 | 0.0001 * * |
| Gram Positive - Actinobacteria | | | | |
| Actinomycetia | Actinomycetales | All | 0 | 1.0000 |
| Actinomycetia | Actinomycetales | Intrasporangiaceae | 0.98 | 0.2730 |
| Actinomycetia | Actinomycetales | Microbacteriaceae | 1.054 | 0.3751 |
| Actinomycetia | Actinomycetales | Nocardioidaceae | 0 | 1.0000 |
| Actinomycetia | All | All | 0 | 1.0000 |
| Gram Positive - All | | | | |
| All | All | All | 0.421 | 0.0325 * |
| Gram Positive - Firmicutes | | | | |
| Bacilli | All | All | 0 | 1.0000 |

#### Supplemental Table 2. Syncom strains

| Strain | Genus | Family | Order | Class | Phylum |
|---|---|---|---|---|---|
| Mixed | | | | | |
| Root70 | Acidovorax | Comamonadaceae | Burkholderiales | Betaproteobacteria | Proteobacteria |
| Root236 | Aeromicrobium | Nocardioidaceae | Propionibacteriales | Actinomycetia | Actinobacteria |
| Root100 | Aminobacter | Phyllobacteriaceae | Rhizobiales | Alphaproteobacteria | Proteobacteria |
| Root239 | Bacillus | Bacillaceae | Bacillales | Bacilli | Firmicutes |
| Root483D1 | Bosea | Bradyrhizobiaceae | Rhizobiales | Alphaproteobacteria | Proteobacteria |
| Root342 | Caulobacter | Caulobacteraceae | Caulobacterales | Alphaproteobacteria | Proteobacteria |
| Root137 | Cellulomonas | Cellulomonadaceae | Actinomycetales | Actinomycetia | Actinobacteria |
| Root1480D1 | Duganella | Oxalobacteraceae | Burkholderiales | Gammaproteobacteria | Proteobacteria |
| Root231 | Ensifer | Rhizobiaceae | Rhizobiales | Alphaproteobacteria | Proteobacteria |
| Root420 | Flavobacterium | Flavobacteriaceae | Flavobacteriales | Bacteroidia | Bacteroidetes |
| Root268 | Hoeflea | Phyllobacteriaceae | Rhizobiales | Alphaproteobacteria | Proteobacteria |
| Root209 | Hydrogenophaga | Comamonadaceae | Burkholderiales | Gammaproteobacteria | Proteobacteria |
| Root107 | Kitasatospora | Streptomycetaceae | Streptomycetales | Actinomycetia | Actinobacteria |
| Root96 | Lysobacter | Xanthomonadaceae | Xanthomonadales | Gammaproteobacteria | Proteobacteria |
| Root133 | Massilia | Oxalobacteraceae | Burkholderiales | Gammaproteobacteria | Proteobacteria |
| Root172 | Mesorhizobium | Phyllobacteriaceae | Rhizobiales | Alphaproteobacteria | Proteobacteria |
| Root180 | Microbacterium | Microbacteriaceae | Actinomycetales | Actinomycetia | Actinobacteria |
| Root1257 | Nocardioides | Nocardioidaceae | Propionibacteriales | Actinomycetia | Actinobacteria |
| Root444D2 | Paenibacillus | Paenibacillaceae | Paenibacillales | Bacilli | Firmicutes |
| Root101 | Phycicoccus | Intrasporangiaceae | Actinomycetales | Actinomycetia | Actinobacteria |
| Root1204 | Rhizobium | Rhizobiaceae | Rhizobiales | Alphaproteobacteria | Proteobacteria |
| Root214 | Sphingopyxis | Sphingomonadaceae | Sphingomonadales | Alphaproteobacteria | Proteobacteria |
| Tolerant | | | | | |
| Root217 | Acidovorax | Comamonadaceae | Burkholderiales | Betaproteobacteria | Proteobacteria |
| Root1280 | Acinetobacter | Moraxellaceae | Pseudomonadales | Gammaproteobacteria | Proteobacteria |
| Root495 | Aeromicrobium | Nocardioidaceae | Propionibacteriales | Actinomycetia | Actinobacteria |
| Root123D2 | Afipia | Bradyrhizobiaceae | Rhizobiales | Alphaproteobacteria | Proteobacteria |
| Soil762 | Arthrobacter | Micrococcaceae | Actinomycetales | Actinomycetia | Actinobacteria |
| Root239 | Bacillus | Bacillaceae | Bacillales | Bacilli | Firmicutes |
| Root565 | Bordetella | Alcaligenaceae | Burkholderiales | Gammaproteobacteria | Proteobacteria |
| Root381 | Bosea | Bradyrhizobiaceae | Rhizobiales | Alphaproteobacteria | Proteobacteria |
| Root1472 | Caulobacter | Caulobacteraceae | Caulobacterales | Alphaproteobacteria | Proteobacteria |
| Root919 | Cupriavidus | Burkholderiaceae | Burkholderiales | Gammaproteobacteria | Proteobacteria |
| Root685 | Devosia | Hyphomicrobiaceae | Rhizobiales | Alphaproteobacteria | Proteobacteria |
| Root1312 | Ensifer | Rhizobiaceae | Rhizobiales | Alphaproteobacteria | Proteobacteria |
| Root1212 | Hoeflea | Phyllobacteriaceae | Rhizobiales | Alphaproteobacteria | Proteobacteria |
| Soil729 | Knoellia | Intrasporangiaceae | Actinomycetales | Actinomycetia | Actinobacteria |
| Root112D2 | Leifsonia | Microbacteriaceae | Actinomycetales | Actinomycetia | Actinobacteria |
| Root133 | Massilia | Oxalobacteraceae | Burkholderiales | Gammaproteobacteria | Proteobacteria |
| Root102 | Mesorhizobium | Phyllobacteriaceae | Rhizobiales | Alphaproteobacteria | Proteobacteria |
| Root1272 | Methylibium | Burkholderiales | Burkholderiales | Gammaproteobacteria | Proteobacteria |
| Root166 | Microbacterium | Microbacteriaceae | Actinomycetales | Actinomycetia | Actinobacteria |
| Root135 | Mycobacterium | Mycobacteriaceae | Mycobacteriales | Actinomycetia | Actinobacteria |
| Root157 | Nitratireductor | Phyllobacteriaceae | Rhizobiales | Alphaproteobacteria | Proteobacteria |
| Root79 | Nocardioides | Nocardioidaceae | Propionibacteriales | Actinomycetia | Actinobacteria |
| Root189 | Noviherbaspirillum | Oxalobacteraceae | Burkholderiales | Gammaproteobacteria | Proteobacteria |
| Root918 | Oerskovia | Cellulomonadaceae | Actinomycetales | Actinomycetia | Actinobacteria |
| Root700 | Phenylobacterium | Caulobacteraceae | Caulobacterales | Alphaproteobacteria | Proteobacteria |

| | | | | | |
|---|---|---|---|---|---|
| Root9 | Pseudomonas | Pseudomonadaceae | Pseudomonadales | Gammaproteobacteria | Proteobacteria |
| Root1203 | Rhizobium | Rhizobiaceae | Rhizobiales | Alphaproteobacteria | Proteobacteria |
| Root561 | Rhodanobacter | Xanthomonadaceae | Xanthomonadales | Gammaproteobacteria | Proteobacteria |
| Root66D1 | Streptomyces | Streptomycetaceae | Streptomycetales | Actinomycetia | Actinobacteria |
| Root85 | Terrabacter | Intrasporangiaceae | Actinomycetales | Actinomycetia | Actinobacteria |
| Soil756 | Tetrasphaera | Intrasporangiaceae | Actinomycetales | Actinomycetia | Actinobacteria |
| Sensitive | | | | | |
| Root70 | Acidovorax | Comamonadaceae | Burkholderiales | Betaproteobacteria | Proteobacteria |
| Root236 | Aeromicrobium | Nocardioidaceae | Propionibacteriales | Actinomycetia | Actinobacteria |
| Root100 | Aminobacter | Phyllobacteriaceae | Rhizobiales | Alphaproteobacteria | Proteobacteria |
| Soil531 | Bacillus | Bacillaceae | Bacillales | Bacilli | Firmicutes |
| Root483D1 | Bosea | Bradyrhizobiaceae | Rhizobiales | Alphaproteobacteria | Proteobacteria |
| Root137 | Cellulomonas | Cellulomonadaceae | Actinomycetales | Actinomycetia | Actinobacteria |
| Root142 | Ensifer | Rhizobiaceae | Rhizobiales | Alphaproteobacteria | Proteobacteria |
| Root935 | Flavobacterium | Flavobacteriaceae | Flavobacteriales | Bacteroidia | Bacteroidetes |
| Root209 | Hydrogenophaga | Comamonadaceae | Burkholderiales | Gammaproteobacteria | Proteobacteria |
| Root983 | Lysobacter | Xanthomonadaceae | Xanthomonadales | Gammaproteobacteria | Proteobacteria |
| Root172 | Mesorhizobium | Phyllobacteriaceae | Rhizobiales | Alphaproteobacteria | Proteobacteria |
| Root61 | Microbacterium | Microbacteriaceae | Actinomycetales | Actinomycetia | Actinobacteria |
| Root240 | Nocardioides | Nocardioidaceae | Propionibacteriales | Actinomycetia | Actinobacteria |
| Soil750 | Paenibacillus | Paenibacillaceae | Paenibacillales | Bacilli | Firmicutes |
| Root1277 | Phenylobacterium | Caulobacteraceae | Caulobacterales | Alphaproteobacteria | Proteobacteria |
| Root563 | Phycicoccus | Intrasporangiaceae | Actinomycetales | Actinomycetia | Actinobacteria |
| Root274 | Rhizobium | Rhizobiaceae | Rhizobiales | Alphaproteobacteria | Proteobacteria |
| Root214 | Sphingopyxis | Sphingomonadaceae | Sphingomonadales | Alphaproteobacteria | Proteobacteria |
| Soil811 | Terrabacter | Intrasporangiaceae | Actinomycetales | Actinomycetia | Actinobacteria |
| Root434 | Variovorax | Comamonadaceae | Burkholderiales | Gammaproteobacteria | Proteobacteria |
| Random | | | | | |
| Root217 | Acidovorax | Comamonadaceae | Burkholderiales | Betaproteobacteria | Proteobacteria |
| Root1280 | Acinetobacter | Moraxellaceae | Pseudomonadales | Gammaproteobacteria | Proteobacteria |
| Root344 | Aeromicrobium | Nocardioidaceae | Propionibacteriales | Actinomycetia | Actinobacteria |
| Root123D2 | Afipia | Bradyrhizobiaceae | Rhizobiales | Alphaproteobacteria | Proteobacteria |
| Root1464 | Agromyces | Microbacteriaceae | Actinomycetales | Actinomycetia | Actinobacteria |
| Soil736 | Arthrobacter | Micrococcaceae | Actinomycetales | Actinomycetia | Actinobacteria |
| Soil531 | Bacillus | Bacillaceae | Bacillales | Bacilli | Firmicutes |
| Root565 | Bordetella | Alcaligenaceae | Burkholderiales | Gammaproteobacteria | Proteobacteria |
| Root381 | Bosea | Bradyrhizobiaceae | Rhizobiales | Alphaproteobacteria | Proteobacteria |
| Root342 | Caulobacter | Caulobacteraceae | Caulobacterales | Alphaproteobacteria | Proteobacteria |
| Root137 | Cellulomonas | Cellulomonadaceae | Actinomycetales | Actinomycetia | Actinobacteria |
| Root919 | Cupriavidus | Burkholderiaceae | Burkholderiales | Gammaproteobacteria | Proteobacteria |
| Root105 | Devosia | Hyphomicrobiaceae | Rhizobiales | Alphaproteobacteria | Proteobacteria |
| Root1480D1 | Duganella | Oxalobacteraceae | Burkholderiales | Gammaproteobacteria | Proteobacteria |
| Root1252 | Ensifer | Rhizobiaceae | Rhizobiales | Alphaproteobacteria | Proteobacteria |
| Root420 | Flavobacterium | Flavobacteriaceae | Flavobacteriales | Bacteroidia | Bacteroidetes |
| Root268 | Hoeflea | Phyllobacteriaceae | Rhizobiales | Alphaproteobacteria | Proteobacteria |
| Soil728 | Janibacter | Intrasporangiaceae | Actinomycetales | Actinomycetia | Actinobacteria |
| Root107 | Kitasatospora | Streptomycetaceae | Streptomycetales | Actinomycetia | Actinobacteria |
| Soil729 | Knoellia | Intrasporangiaceae | Actinomycetales | Actinomycetia | Actinobacteria |
| Root1293 | Leifsonia | Microbacteriaceae | Actinomycetales | Actinomycetia | Actinobacteria |
| Root604 | Lysobacter | Xanthomonadaceae | Xanthomonadales | Gammaproteobacteria | Proteobacteria |
| Root133 | Massilia | Oxalobacteraceae | Burkholderiales | Gammaproteobacteria | Proteobacteria |
| Root1471 | Mesorhizobium | Phyllobacteriaceae | Rhizobiales | Alphaproteobacteria | Proteobacteria |

| | | | | | |
|---|---|---|---|---|---|
| Root1272 | Methylibium | Burkholderiales | Burkholderiales | Gammaproteobacteria | Proteobacteria |
| Root322 | Microbacterium | Microbacteriaceae | Actinomycetales | Actinomycetia | Actinobacteria |
| Soil538 | Mycobacterium | Mycobacteriaceae | Mycobacteriales | Actinomycetia | Actinobacteria |
| Root157 | Nitratireductor | Phyllobacteriaceae | Rhizobiales | Alphaproteobacteria | Proteobacteria |
| Root136 | Nocardia | Nocardiaceae | Mycobacteriales | Actinomycetia | Actinobacteria |
| Root1257 | Nocardioides | Nocardioidaceae | Propionibacteriales | Actinomycetia | Actinobacteria |
| Root189 | Noviherbaspirillum | Oxalobacteraceae | Burkholderiales | Gammaproteobacteria | Proteobacteria |
| Root22 | Oerskovia | Cellulomonadaceae | Actinomycetales | Actinomycetia | Actinobacteria |
| Root52 | Paenibacillus | Paenibacillaceae | Paenibacillales | Bacilli | Firmicutes |
| Root700 | Phenylobacterium | Caulobacteraceae | Caulobacterales | Alphaproteobacteria | Proteobacteria |
| Root563 | Phycicoccus | Intrasporangiaceae | Actinomycetales | Actinomycetia | Actinobacteria |
| Root329 | Pseudomonas | Pseudomonadaceae | Pseudomonadales | Gammaproteobacteria | Proteobacteria |
| Root65 | Pseudoxanthomonas | Xanthomonadaceae | Xanthomonadales | Gammaproteobacteria | Proteobacteria |
| Root29 | Rhizobacter | Pseudomonadaceae | Burkholderiales | Gammaproteobacteria | Proteobacteria |
| Root708 | Rhizobium | Rhizobiaceae | Rhizobiales | Alphaproteobacteria | Proteobacteria |
| Soil772 | Rhodanobacter | Xanthomonadaceae | Xanthomonadales | Gammaproteobacteria | Proteobacteria |
| Root1294 | Sphingomonas | Sphingomonadaceae | Sphingomonadales | Alphaproteobacteria | Proteobacteria |
| Root214 | Sphingopyxis | Sphingomonadaceae | Sphingomonadales | Alphaproteobacteria | Proteobacteria |
| Root1310 | Streptomyces | Streptomycetaceae | Streptomycetales | Actinomycetia | Actinobacteria |
| Soil811 | Terrabacter | Intrasporangiaceae | Actinomycetales | Actinomycetia | Actinobacteria |
| Soil756 | Tetrasphaera | Intrasporangiaceae | Actinomycetales | Actinomycetia | Actinobacteria |
| Root411 | Variovorax | Comamonadaceae | Burkholderiales | Gammaproteobacteria | Proteobacteria |
| Root332 | Yonghaparkia | Microbacteriaceae | Actinomycetales | Actinomycetia | Actinobacteria |

#### Supplemental Table 3. PERMANOVA

| term | Df | SumsOfSqs | MeanSqs | F.Model | R2 | p.value |
|---|---|---|---|---|---|---|
| Random | | | | | | |
| Timepoint | 3 | 1.1953 | 0.3984 | 14.7627 | 0.3100 | 0.001 * * |
| Treatment | 2 | 1.5522 | 0.7761 | 28.7561 | 0.4026 | 0.001 * * |
| Timepoint:Treatment | 6 | 0.1363 | 0.0227 | 0.8420 | 0.0354 | 0.594 |
| Residuals | 36 | 0.9716 | 0.0270 | NA | 0.2520 | NA |
| Total | 47 | 3.8555 | NA | NA | 1.0000 | NA |
| Tolerant | | | | | | |
| Timepoint | 3 | 0.9527 | 0.3176 | 10.9268 | 0.3293 | 0.001 * * |
| Treatment | 2 | 0.6252 | 0.3126 | 10.7561 | 0.2161 | 0.001 * * |
| Timepoint:Treatment | 6 | 0.2687 | 0.0448 | 1.5410 | 0.0929 | 0.098 |
| Residuals | 36 | 1.0463 | 0.0291 | NA | 0.3617 | NA |
| Total | 47 | 2.8929 | NA | NA | 1.0000 | NA |
| Mixed | | | | | | |
| Timepoint | 3 | 0.1909 | 0.0636 | 1.2030 | 0.0199 | 0.296 |
| Treatment | 2 | 7.2278 | 3.6139 | 68.3271 | 0.7544 | 0.001 * * |
| Timepoint:Treatment | 6 | 0.2575 | 0.0429 | 0.8115 | 0.0269 | 0.621 |
| Residuals | 36 | 1.9041 | 0.0529 | NA | 0.1987 | NA |
| Total | 47 | 9.5803 | NA | NA | 1.0000 | NA |
| Sensitive | | | | | | |
| Timepoint | 3 | 0.0395 | 0.0132 | 1.9094 | 0.0631 | 0.130 |
| Treatment | 2 | 0.2605 | 0.1303 | 18.9085 | 0.4166 | 0.001 * * |
| Timepoint:Treatment | 6 | 0.0774 | 0.0129 | 1.8728 | 0.1238 | 0.081 |
| Residuals | 36 | 0.2480 | 0.0069 | NA | 0.3966 | NA |
| Total | 47 | 0.6254 | NA | NA | 1.0000 | NA |

#### Supplemental Table 4. PERMANOVA comparing only two treatments

| term | Df | SumsOfSqs | MeanSqs | F.Model | R2 | p.value |
|---|---|---|---|---|---|---|
| Random - APO | | | | | | |
| Treatment | 1 | 1.0477 | 1.0477 | 18.1201 | 0.3766 | 0.001 * * |
| Residuals | 30 | 1.7346 | 0.0578 | NA | 0.6234 | NA |
| Total | 31 | 2.7823 | NA | NA | 1.0000 | NA |
| Random - BOA | | | | | | |
| Treatment | 1 | 0.0412 | 0.0412 | 0.8416 | 0.0273 | 0.435 |
| Residuals | 30 | 1.4680 | 0.0489 | NA | 0.9727 | NA |
| Total | 31 | 1.5092 | NA | NA | 1.0000 | NA |
| Tolerant - APO | | | | | | |
| Treatment | 1 | 0.4742 | 0.4742 | 8.3739 | 0.2182 | 0.001 * * |
| Residuals | 30 | 1.6988 | 0.0566 | NA | 0.7818 | NA |
| Total | 31 | 2.1730 | NA | NA | 1.0000 | NA |
| Tolerant - BOA | | | | | | |
| Treatment | 1 | 0.0617 | 0.0617 | 1.5378 | 0.0488 | 0.197 |
| Residuals | 30 | 1.2028 | 0.0401 | NA | 0.9512 | NA |
| Total | 31 | 1.2645 | NA | NA | 1.0000 | NA |
| Mixed - APO | | | | | | |
| Treatment | 1 | 5.1799 | 5.1799 | 70.8788 | 0.7026 | 0.001 * * |
| Residuals | 30 | 2.1924 | 0.0731 | NA | 0.2974 | NA |
| Total | 31 | 7.3724 | NA | NA | 1.0000 | NA |
| Mixed - BOA | | | | | | |
| Treatment | 1 | 0.0577 | 0.0577 | 4.6442 | 0.1341 | 0.020 * |
| Residuals | 30 | 0.3726 | 0.0124 | NA | 0.8659 | NA |
| Total | 31 | 0.4303 | NA | NA | 1.0000 | NA |
| Sensitive - APO | | | | | | |
| Treatment | 1 | 0.0229 | 0.0229 | 20.8918 | 0.4105 | 0.001 * * |
| Residuals | 30 | 0.0329 | 0.0011 | NA | 0.5895 | NA |
| Total | 31 | 0.0559 | NA | NA | 1.0000 | NA |
| Sensitive - BOA | | | | | | |
| Treatment | 1 | 0.1120 | 0.1120 | 10.3315 | 0.2562 | 0.003 * * |
| Residuals | 30 | 0.3251 | 0.0108 | NA | 0.7438 | NA |
| Total | 31 | 0.4370 | NA | NA | 1.0000 | NA |

#### Supplemental Table 5. ANOVA: Observed taxa

| term | df | sumsq | meansq | statistic | p.value |
|---|---|---|---|---|---|
| Random | | | | | |
| Treatment | 2 | 0.05101594 | 0.025507971 | 4.1016798 | 2.484563e-02 * |
| Timepoint | 3 | 0.25046221 | 0.083487403 | 13.4247681 | 4.907087e-06 * * |
| Treatment:Timepoint | 6 | 0.01463861 | 0.002439768 | 0.3923145 | 8.790628e-01 |
| Residuals | 36 | 0.22388070 | 0.006218908 | NA | NA |
| Tolerant | | | | | |
| Treatment | 2 | 0.21585880 | 0.107929398 | 13.1512316 | 5.156453e-05 * * |
| Timepoint | 3 | 0.32902545 | 0.109675150 | 13.3639521 | 5.118265e-06 * * |
| Treatment:Timepoint | 6 | 0.03393162 | 0.005655270 | 0.6890965 | 6.596909e-01 |
| Residuals | 36 | 0.29544444 | 0.008206790 | NA | NA |
| Mixed | | | | | |
| Treatment | 2 | 2.32954036 | 1.164770180 | 41.3453120 | 4.720082e-10 * * |
| Timepoint | 3 | 0.35051038 | 0.116836792 | 4.1473019 | 1.268430e-02 * |
| Treatment:Timepoint | 6 | 0.31516138 | 0.052526897 | 1.8645231 | 1.140745e-01 |
| Residuals | 36 | 1.01418334 | 0.028171759 | NA | NA |
| Sensitive | | | | | |
| Treatment | 2 | 0.46631836 | 0.233159182 | 30.3635644 | 1.877482e-08 * * |
| Timepoint | 3 | 0.07701343 | 0.025671142 | 3.3430696 | 2.974336e-02 * |
| Treatment:Timepoint | 6 | 0.09728286 | 0.016213809 | 2.1114718 | 7.590701e-02 |
| Residuals | 36 | 0.27644088 | 0.007678913 | NA | NA |

**Supplemental Table 6. Observed diversity means and pairwise comparisons**

| Treatment | estimate | 95% CI | Group[1] |
|---|---|---|---|
| Random - 24h | | | |
| BOA | 1.148 | [1.068 - 1.228] | a |
| APO | 1.216 | [1.136 - 1.296] | a |
| Control | 1.224 | [1.144 - 1.304] | a |
| Random - 48h | | | |
| BOA | 1.221 | [1.141 - 1.301] | a |
| APO | 1.282 | [1.202 - 1.362] | a |
| Control | 1.297 | [1.217 - 1.377] | a |
| Random - 72h | | | |
| APO | 1.329 | [1.249 - 1.409] | a |
| BOA | 1.349 | [1.269 - 1.429] | a |
| Control | 1.435 | [1.355 - 1.515] | a |
| Random - 96h | | | |
| BOA | 1.328 | [1.248 - 1.408] | a |
| APO | 1.351 | [1.271 - 1.431] | a |
| Control | 1.407 | [1.327 - 1.487] | a |
| Tolerant - 24h | | | |
| BOA | 0.964 | [0.872 - 1.056] | a |
| Control | 0.994 | [0.902 - 1.086] | ab |
| APO | 1.125 | [1.033 - 1.217] | b |
| Tolerant - 48h | | | |
| Control | 1.055 | [0.963 - 1.147] | a |
| BOA | 1.145 | [1.053 - 1.237] | ab |
| APO | 1.275 | [1.183 - 1.367] | b |
| Tolerant - 72h | | | |
| Control | 1.128 | [1.036 - 1.22] | a |
| BOA | 1.165 | [1.073 - 1.257] | ab |
| APO | 1.315 | [1.223 - 1.407] | b |
| Tolerant - 96h | | | |
| Control | 1.208 | [1.117 - 1.3] | a |
| BOA | 1.247 | [1.155 - 1.339] | a |
| APO | 1.295 | [1.203 - 1.387] | a |
| Mixed - 24h | | | |
| BOA | 0.539 | [0.368 - 0.709] | a |
| Control | 0.666 | [0.496 - 0.836] | ab |
| APO | 0.907 | [0.737 - 1.078] | b |
| Mixed - 48h | | | |
| BOA | 0.228 | [0.057 - 0.398] | a |
| Control | 0.344 | [0.174 - 0.514] | a |
| APO | 1.027 | [0.856 - 1.197] | b |
| Mixed - 72h | | | |
| BOA | 0.250 | [0.08 - 0.42] | a |

| | | | |
|---|---|---|---|
| Control | 0.423 | [0.253 - 0.593] | a |
| APO | 0.739 | [0.569 - 0.909] | b |
| Mixed - 96h | | | |
| BOA | 0.385 | [0.215 - 0.555] | a |
| Control | 0.532 | [0.361 - 0.702] | ab |
| APO | 0.814 | [0.644 - 0.984] | b |
| Sensitive - 24h | | | |
| APO | 0.041 | [-0.048 - 0.13] | a |
| Control | 0.210 | [0.121 - 0.299] | b |
| BOA | 0.220 | [0.131 - 0.309] | b |
| Sensitive - 48h | | | |
| APO | 0.044 | [-0.045 - 0.133] | a |
| Control | 0.144 | [0.055 - 0.233] | a |
| BOA | 0.377 | [0.289 - 0.466] | b |
| Sensitive - 72h | | | |
| APO | 0.024 | [-0.065 - 0.113] | a |
| Control | 0.057 | [-0.032 - 0.146] | a |
| BOA | 0.308 | [0.22 - 0.397] | b |
| Sensitive - 96h | | | |
| APO | 0.021 | [-0.068 - 0.109] | a |
| Control | 0.043 | [-0.045 - 0.132] | ab |
| BOA | 0.174 | [0.085 - 0.262] | b |

[1]Different letters indicate significant differences within one Syncom / Timepoint combination (alpha = 0.05), Tukey adjusted

#### Supplemental Table 7. ANOVA: Shannon Diversity

| term | df | sumsq | meansq | statistic | p.value |
|---|---|---|---|---|---|
| Random | | | | | |
| Treatment | 2 | 0.04965670 | 0.024828351 | 3.9818448 | 2.740000e-02 * |
| Timepoint | 3 | 0.25834405 | 0.086114684 | 13.8106353 | 3.764333e-06 * * |
| Treatment:Timepoint | 6 | 0.02053069 | 0.003421782 | 0.5487681 | 7.675710e-01 |
| Residuals | 36 | 0.22447400 | 0.006235389 | NA | NA |
| Tolerant | | | | | |
| Treatment | 2 | 0.20619734 | 0.103098670 | 11.8582100 | 1.106005e-04 * * |
| Timepoint | 3 | 0.31193275 | 0.103977584 | 11.9593009 | 1.392757e-05 * * |
| Treatment:Timepoint | 6 | 0.03231331 | 0.005385552 | 0.6194357 | 7.133662e-01 |
| Residuals | 36 | 0.31299430 | 0.008694286 | NA | NA |
| Mixed | | | | | |
| Treatment | 2 | 2.20186153 | 1.100930766 | 37.4317591 | 1.611570e-09 * * |
| Timepoint | 3 | 0.31214721 | 0.104049071 | 3.5376791 | 2.413246e-02 * |
| Treatment:Timepoint | 6 | 0.31328626 | 0.052214376 | 1.7752942 | 1.320402e-01 |
| Residuals | 36 | 1.05882033 | 0.029411676 | NA | NA |
| Sensitive | | | | | |
| Treatment | 2 | 0.64164868 | 0.320824339 | 33.1316491 | 6.894276e-09 * * |
| Timepoint | 3 | 0.09113328 | 0.030377760 | 3.1371226 | 3.717840e-02 * |
| Treatment:Timepoint | 6 | 0.12710073 | 0.021183455 | 2.1876233 | 6.692054e-02 |
| Residuals | 36 | 0.34859950 | 0.009683319 | NA | NA |

#### Supplemental Table 8. Shannon diversity means and pairwise comparisons

| Treatment | estimate | 95% CI | Group[1] |
|---|---|---|---|
| Random - 24h | | | |
| BOA | 1.130 | [1.05 - 1.21] | a |
| Control | 1.224 | [1.144 - 1.305] | a |
| APO | 1.225 | [1.145 - 1.305] | a |
| Random - 48h | | | |
| BOA | 1.235 | [1.155 - 1.315] | a |
| APO | 1.282 | [1.202 - 1.362] | a |
| Control | 1.296 | [1.216 - 1.376] | a |
| Random - 72h | | | |
| APO | 1.329 | [1.249 - 1.409] | a |
| BOA | 1.354 | [1.274 - 1.435] | a |
| Control | 1.437 | [1.357 - 1.517] | a |
| Random - 96h | | | |
| BOA | 1.330 | [1.25 - 1.41] | a |
| APO | 1.349 | [1.269 - 1.429] | a |
| Control | 1.406 | [1.326 - 1.486] | a |
| Tolerant - 24h | | | |
| BOA | 0.995 | [0.9 - 1.09] | a |
| Control | 1.023 | [0.928 - 1.117] | a |
| APO | 1.142 | [1.047 - 1.236] | a |
| Tolerant - 48h | | | |
| Control | 1.071 | [0.976 - 1.166] | a |
| BOA | 1.162 | [1.067 - 1.256] | ab |
| APO | 1.290 | [1.195 - 1.384] | b |
| Tolerant - 72h | | | |
| Control | 1.148 | [1.053 - 1.242] | a |
| BOA | 1.182 | [1.087 - 1.276] | ab |
| APO | 1.333 | [1.239 - 1.428] | b |
| Tolerant - 96h | | | |
| Control | 1.231 | [1.137 - 1.326] | a |
| BOA | 1.263 | [1.169 - 1.358] | a |
| APO | 1.317 | [1.223 - 1.412] | a |
| Mixed - 24h | | | |
| BOA | 0.517 | [0.343 - 0.691] | a |
| Control | 0.651 | [0.477 - 0.825] | ab |
| APO | 0.917 | [0.743 - 1.091] | b |
| Mixed - 48h | | | |
| BOA | 0.233 | [0.059 - 0.407] | a |
| Control | 0.353 | [0.179 - 0.527] | a |
| APO | 1.021 | [0.847 - 1.195] | b |
| Mixed - 72h | | | |
| BOA | 0.258 | [0.084 - 0.432] | a |

| | | | |
|---|---|---|---|
| Control | 0.443 | [0.269 - 0.617] | ab |
| APO | 0.723 | [0.549 - 0.897] | b |
| Mixed - 96h | | | |
| BOA | 0.402 | [0.228 - 0.576] | a |
| Control | 0.555 | [0.381 - 0.729] | ab |
| APO | 0.789 | [0.615 - 0.962] | b |
| Sensitive - 24h | | | |
| APO | 0.043 | [-0.057 - 0.143] | a |
| Control | 0.234 | [0.134 - 0.334] | b |
| BOA | 0.247 | [0.147 - 0.347] | b |
| Sensitive - 48h | | | |
| APO | 0.051 | [-0.049 - 0.151] | a |
| Control | 0.172 | [0.072 - 0.272] | a |
| BOA | 0.439 | [0.339 - 0.539] | b |
| Sensitive - 72h | | | |
| APO | 0.032 | [-0.068 - 0.131] | a |
| Control | 0.073 | [-0.027 - 0.173] | a |
| BOA | 0.368 | [0.268 - 0.467] | b |
| Sensitive - 96h | | | |
| APO | 0.027 | [-0.073 - 0.127] | a |
| Control | 0.056 | [-0.044 - 0.156] | ab |
| BOA | 0.214 | [0.114 - 0.314] | b |

[1]Different letters indicate significant differences within one Syncom / Timepoint combination (alpha = 0.05), Tukey adjusted

**Supplemental Table 9. Correlation between change in relative abundance and change in growth**

| | r2 | Treatment |
|---|---|---|
| log2FC vs absolute AUC | | |
| | 0.01983379 | APO |
| | -0.01052433 | BOA |
| log2FC vs relative AUC | | |
| | 0.01071874 | APO |
| | 0.01237386 | BOA |

### Figures

#### Supplemental Figure 1

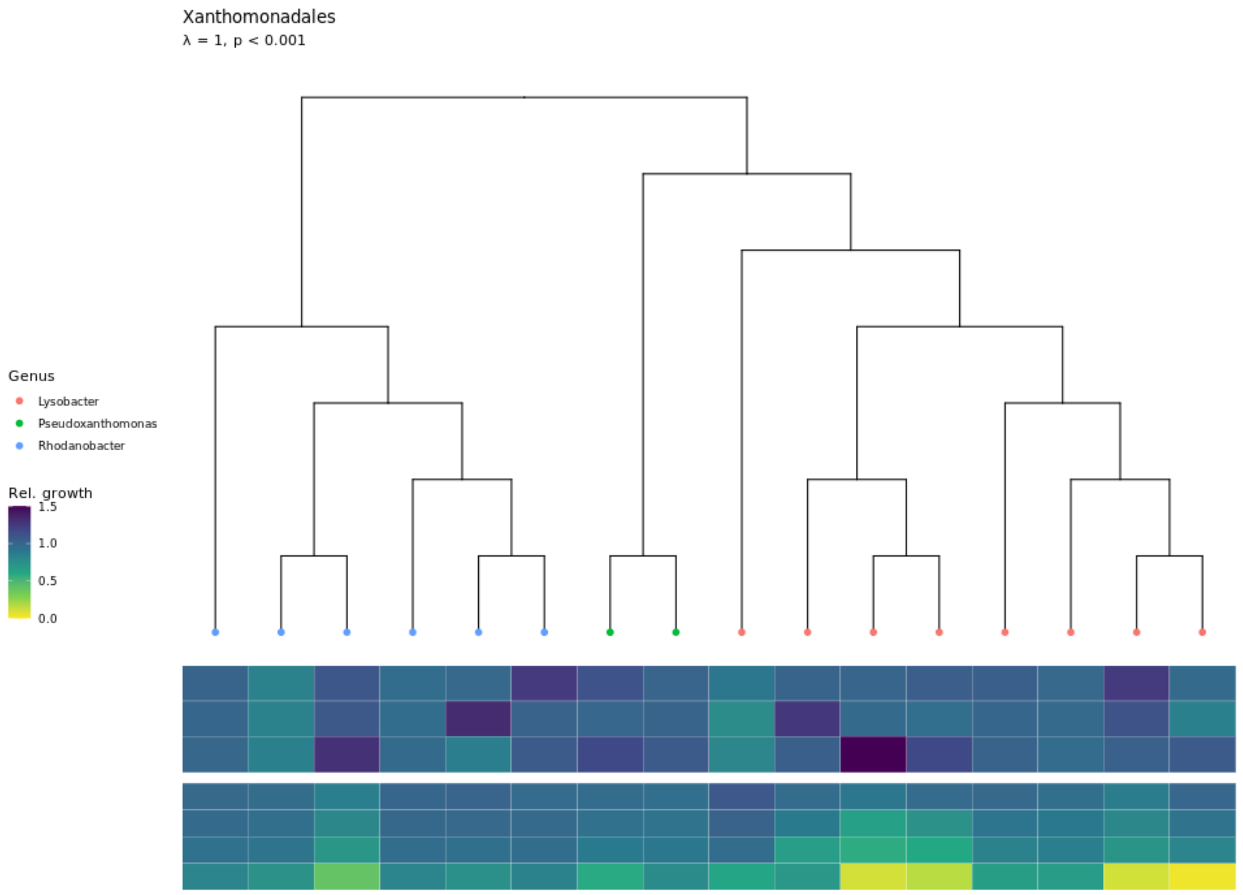

**Supplemental Figure 1. Bx sensitivity of Xanthomonadales.** Genera belonging to the Order Xanthomonadales are plotted. The tree was constructed from concatenated sequences (see methods). The upper 3 rows of the heatmap display relative growth in 10µM, 50µM and 100µM BOA (top to bottom), the lower four rows of the heatmap relative growth in medium supplemented with 1µM, 5µM, 10µM and 50µM APO.

#### Supplemental Figure 2

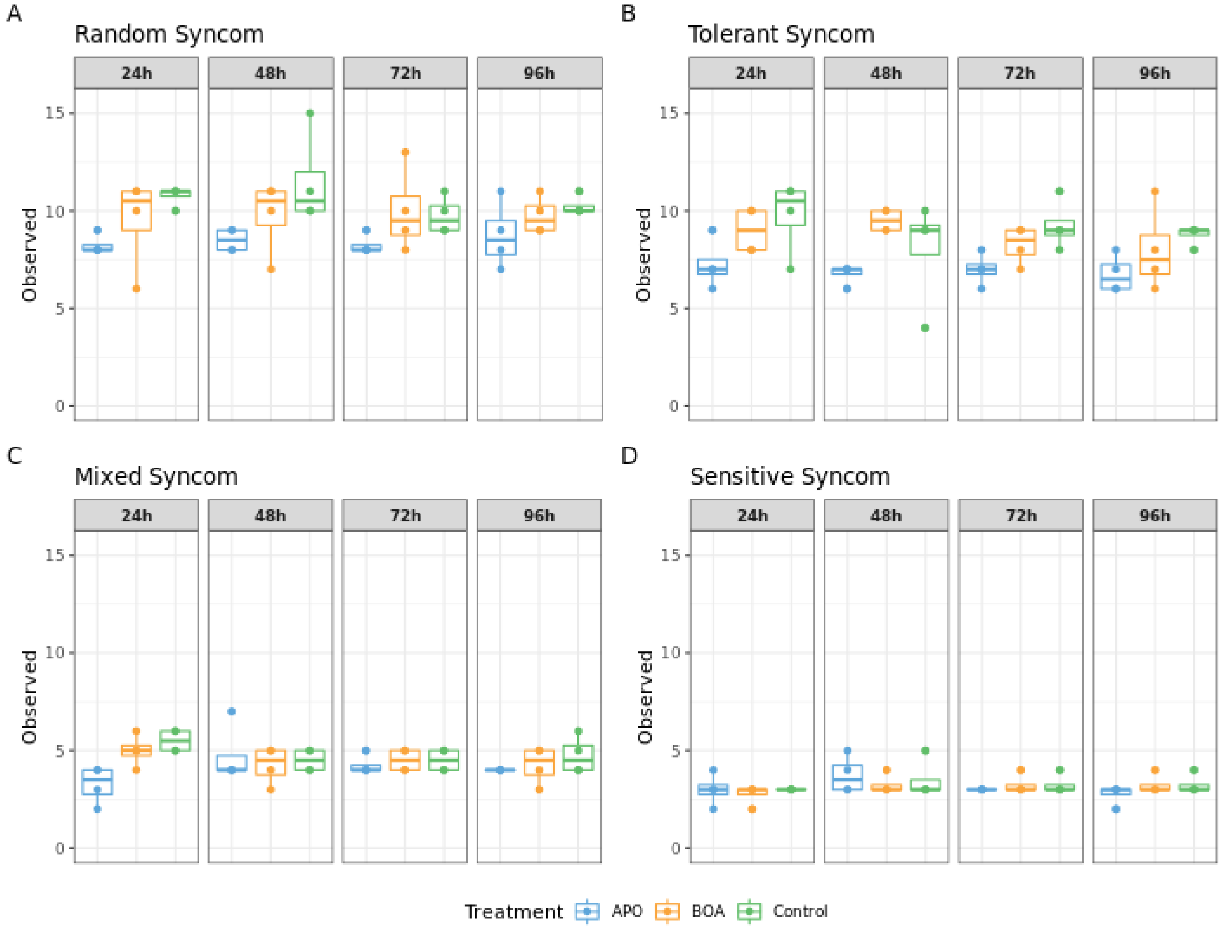

**Supplemental Figure 2. Observed alpha diversity.** The number of observed bacterial genera of the samples belonging to the four different syncoms (**A-D**) is shown for each time point (facet) and treatment (color). Each sample is plotted as a dot, and box-and-whisker plots show the summary statistics for each treatment / time point combination.

#### Supplemental Figure 3

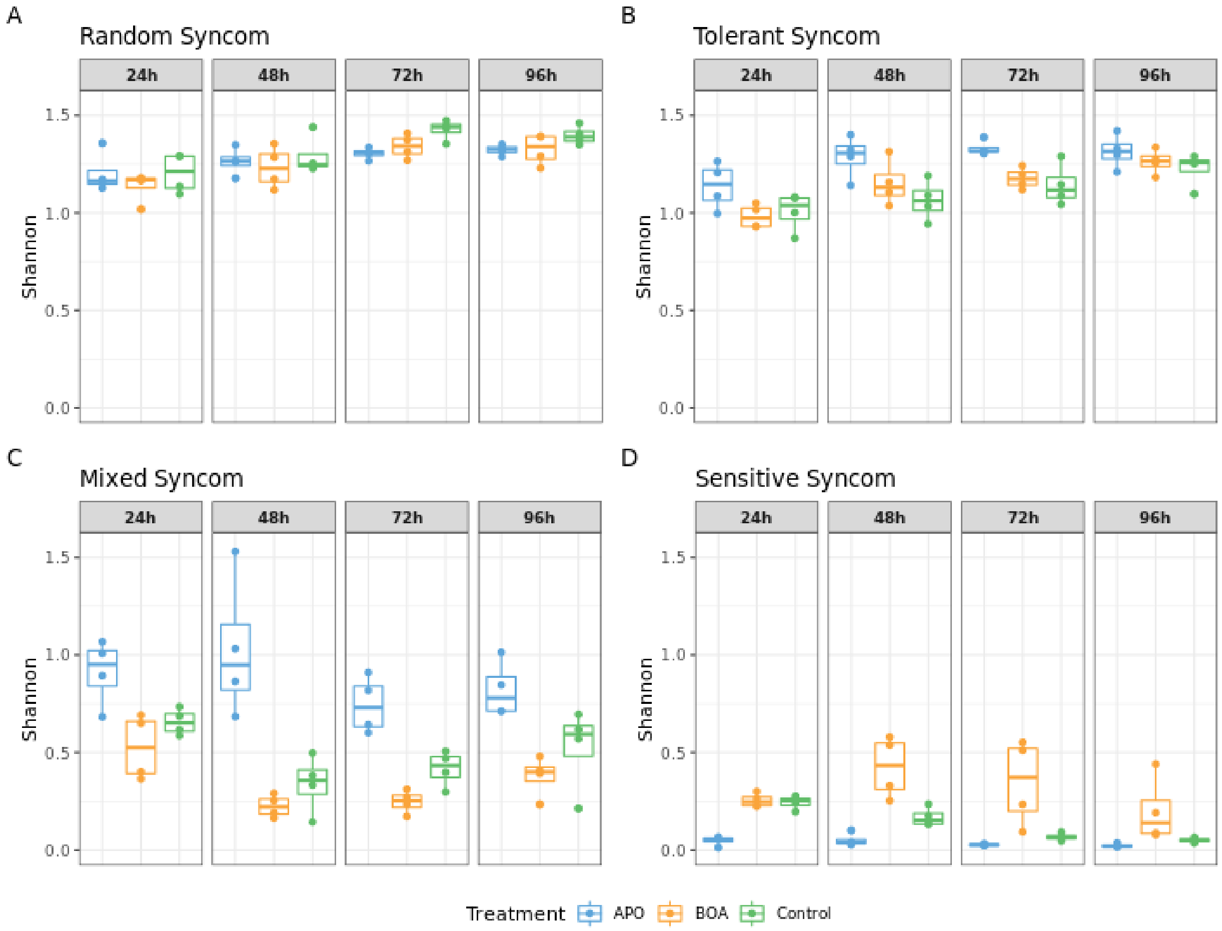

**Supplemental Figure 3. Shannon diversity indices of the synthetic communities.** The Shannon indices of the samples belonging to the four different syncoms (*A*-*D*) are shown for each time point (facet) and treatment (color). Each sample is plotted as a dot, and box-and-whisker plots show the summary statistics for each treatment / time point combination.

### Supplemental Figure 4

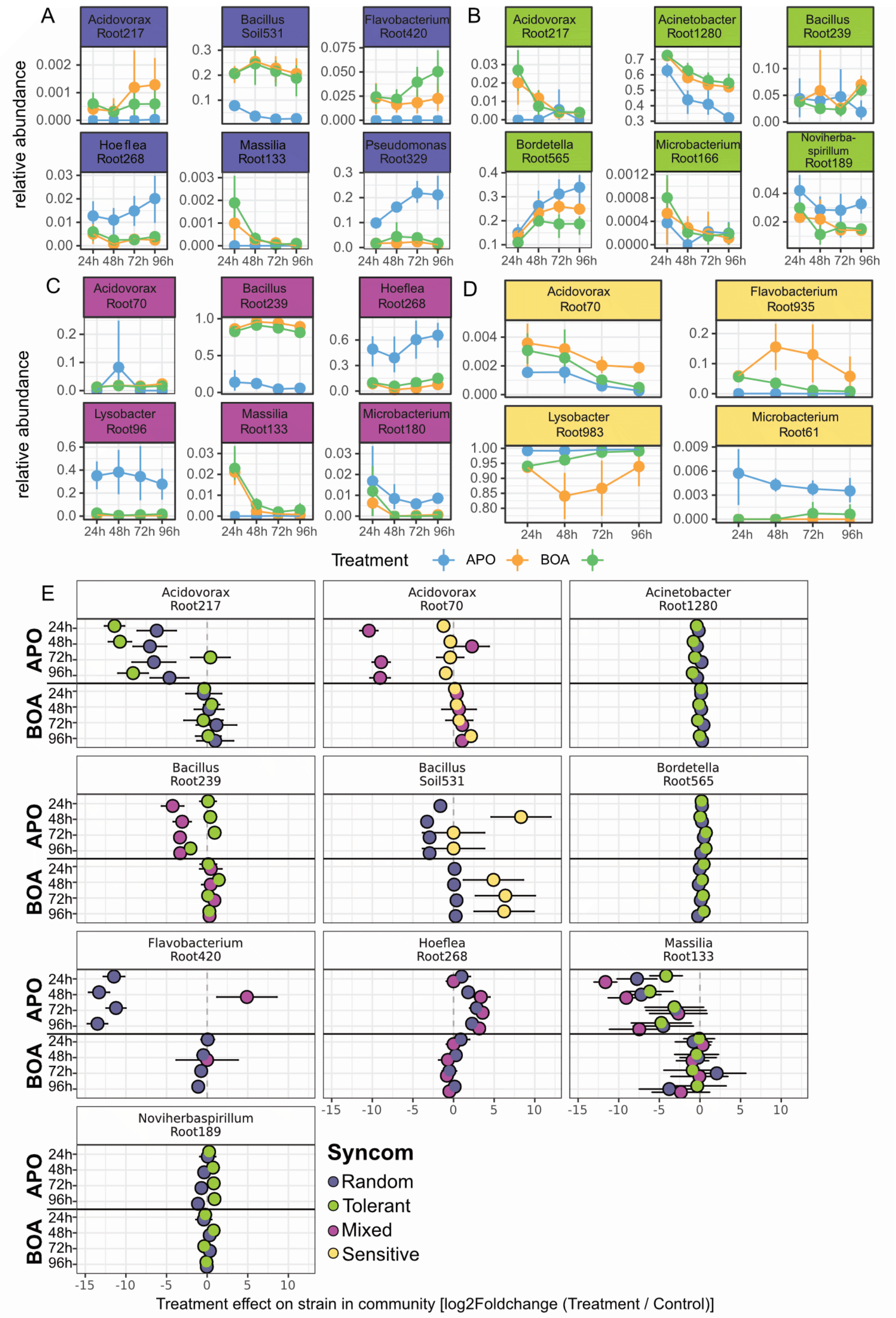

**Supplemental Figure 4. Changes in relative abundance by community. A-D** The relative abundance over time for different isolates that showed signficant differences in relative abundance at at least one timepoint (compared to control). Relative abundance is shown per treatment and timepoint, errorbars indicate 95% CI. **E** Changes in relative abundance per treatment and timepoint for isolates that were found in more than one community. Errorbars indicate s.e. of the log2Foldchange.
